# Widespread gene regulator Psu inhibits transcription termination factor ρ by forced hyper-oligomerization

**DOI:** 10.1101/2023.06.22.546067

**Authors:** Daniela Gjorgjevikj, Naveen Kumar, Bing Wang, Tarek Hilal, Nelly Said, Bernhard Loll, Irina Artsimovitch, Ranjan Sen, Markus C. Wahl

## Abstract

Many bacteriophages modulate the host transcription machinery for efficient expression of their own genomes. Phage P4 polarity suppression protein, Psu, is a building block of the viral capsid and inhibits the hexameric transcription termination factor, ρ, by presently unknown mechanisms. We elucidated cryogenic electron microscopy structures of ρ-Psu complexes, showing that Psu dimers laterally clamp two inactive, open ρ rings and promote their expansion to higher-oligomeric states. Systematic ATPase, nucleotide binding and nucleic acid binding studies revealed that Psu hinders ρ ring closure and traps nucleotides in their binding pockets on ρ. Structure-guided mutagenesis in combination with growth, pull-down and termination assays further delineated the functional ρ-Psu interfaces. Bioinformatic analyses suggested that, in addition to guarding its own genome against ρ, Psu enables expression of diverse phage-defense systems commonly found in P4-like mobile genetic elements across bacteria. Thus, Psu is a widespread gene regulator that inhibits ρ *via* forced hyper-oligomerization.

## Main

More than 90 % of bacterial species encode ρ, a hexameric, ring-shaped RNA-dependent NTPase.^1^ ρ is long known as a mediator of transcription termination^2^ that defines the ends of 20-30 % of transcription units in *Escherichia coli*^3^. ρ also mediates attenuation in 5’-untranslated regions (UTRs)^4^, limits antisense transcription^5, 6^, and safeguards bacterial genomes by restricting R-loops^7^. A major function of ρ is the silencing of horizontally acquired genes and invading foreign DNA^8^. Thus, phages that utilize the bacterial transcription machinery must counteract ρ activity to efficiently express their genomes. While lambdoid phages implement protein- or RNA-based anti termination mechanisms to render the host RNA polymerase (RNAP) termination-resistant^9–15^, the enterobacterial pirate phage, P4, has acquired a unique anti-termination system that acts directly upon ρ. Its polarity suppression (Psu) protein is a building block of its capsid^16^, but can also inhibit ρ^17, 18^, as first noted by its suppression of ρ-dependent polarity^19^. The precise mode of Psu dependent ρ inhibition has so far remained elusive.

A ρ protomer comprises N-terminal and C-terminal domains (NTD and CTD) connected by a flexible hinge region.^20, 21^ The NTD harbors a primary RNA-binding site (PBS).^20, 21^ The CTDs together form a secondary RNA-binding site (SBS) in the central pore of the hexamer and six nucleotide-binding pockets between the subunits.^20, 21^ The ρ hexamer adopts an NTPase-inactive, washer-like, open-ring conformation until NTPs (preferentially ATP) are engaged and RNA is bound at the SBS, leading to ring closure and RNA entrapment inside the ring.^22^ The SBS and the sixth nucleotide-binding site are fully formed only in the closed ρ ring. After RNA engagement at the PBS and the SBS, ρ can act as an NTP hydrolysis-driven 5’-to-3’ RNA-translocase and helicase.^20, 21^

Extensive research over decades suggested two mechanisms of ρ termination. In a tethered tracking model, ρ in the open conformation engages a C-rich, secondary-structure-deficient ρ-utilization (*rut*) site on the nascent RNA (a ρ-dependent terminator) *via* its PBSes.^20, 21^ Subsequently, downstream RNA can enter the SBS, followed by ring closure and NTPase-driven 5’-to-3’ translocation on the transcript, during which the PBSes are thought to remain bound to *rut*.^20, 21^ When RNAP reaches a pause signal, ρ catches up and, using its strong motor activity, disassembles the elongation complex (EC).^23–25^ In an alternative, allosteric model, the NTDs of an open ρ ring engage ECs *via* direct contacts to RNAP and bound general transcription factors (TFs) NusA and NusG^26–28^ and ρ traffics with the EC^29^. Once the EC pauses, ρ can stepwise inactivate RNAP without resorting to its motor activity.^26–28^ Recent single-molecule spectroscopic studies have suggested that, at least *in vitro*, both mechanisms may be at work to a different extent depending on the transcription unit or termination scenario.^30, 31^

Here, we present cryogenic electron microscopy (cryoEM)/single-particle analysis (SPA) based structures of ρ-Psu complexes, defining the precise ρ-Psu interfaces and revealing a unique inhibitory mechanism. Multiple Psu dimers staple two open ρ complexes together and promote the formation of higher-order ρ oligomers. Psu subunits bind between two neighboring ρ CTDs across the nucleotide-binding sites. Our systematic ATPase, nucleotide-binding and nucleic-acid-binding studies are fully consistent with Psu blocking the ρ nucleotide-binding sites and stabilizing ρ in an open conformation, while leaving ρ PBSes unobstructed. Furthermore, we delineate key residues on Psu and ρ required for ρ inhibition *in vitro* and *in vivo*. We also find Psu proteins encoded in the neighborhood of known and putative phage defense genes in many bacterial genomes. Based on our results, we propose that Psu inhibits ρ-dependent transcription termination *via* an unconventional molecular mechanism of forced hyper-oligomerization, which may well be employed by diverse bacteria to regulate the expression of phage defense systems.

## Results

### Psu dimers bridge and promote the expansion of two open ρ hexamers

Nucleotide-free, wild-type (wt) ρ did not form a stable complex with Psu in analytical size exclusion chromatography (SEC; Extended Data Fig. 1a). However, in the presence of ADP-BeF3 or ATPγS, ρ^wt^ and Psu partially co-eluted earlier from the SEC column than either protein alone, suggesting that dynamic ρ^wt^-Psu complexes are stabilized by the nucleotides (Extended Data Fig. 1a). To elucidate the structural basis of ρ^wt^-ATPγS-Psu complex formation, we subjected corresponding mixtures to cryoEM/SPA (Extended Data Fig. 2 and 3; Supplementary Table 1). Multi-particle 3D refinement with ∼750,000 particle images yielded several cryoEM reconstructions (Extended Data Fig. 3). The assemblies each contain two open ρ^wt^ complexes, but with varying numbers of subunits, and different numbers of Psu dimers that bridge the ρ^wt^ complexes. As exemplars, we further refined and analyzed one of the smallest and the largest assembly imaged.

**Fig. 1:**
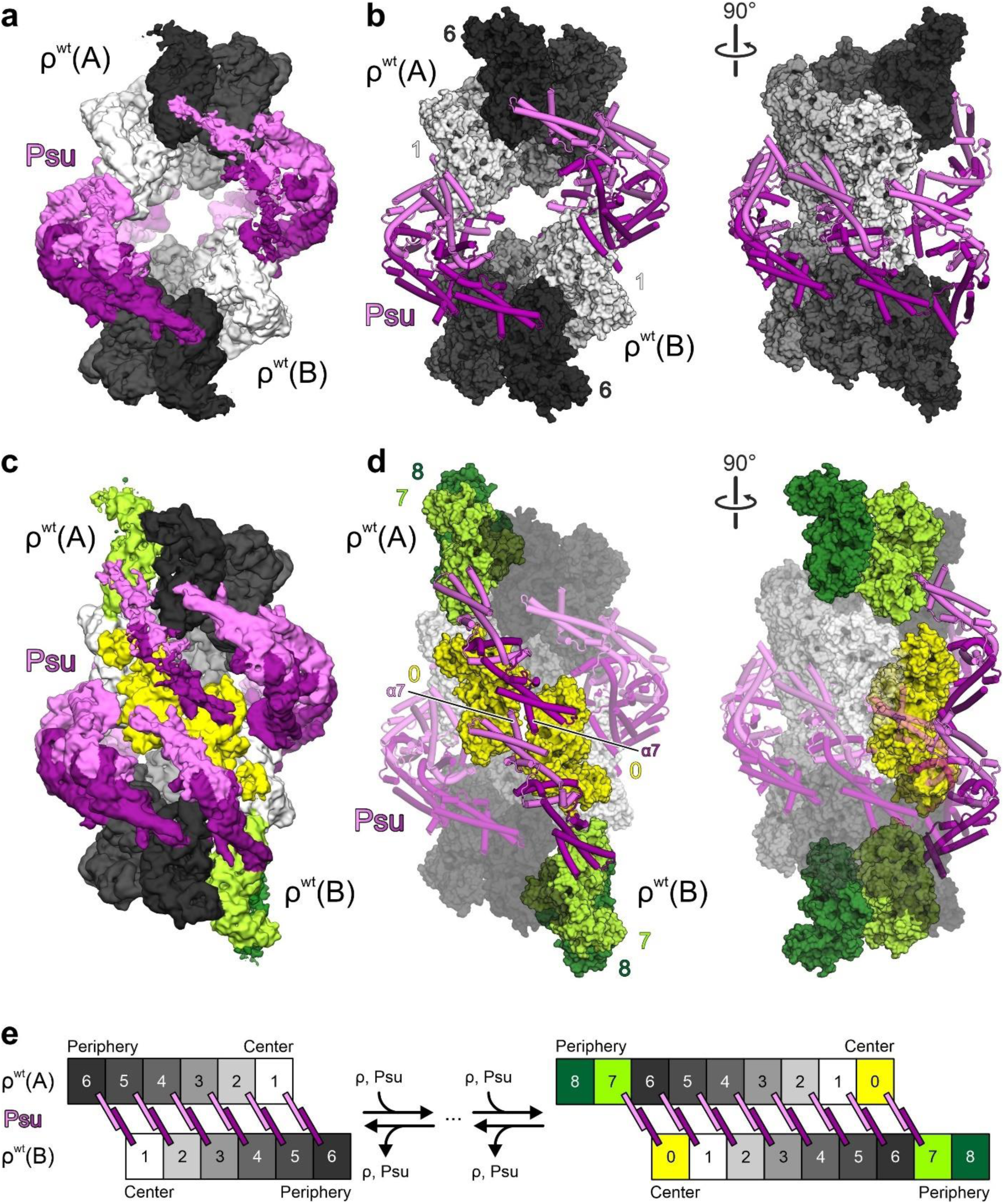
Structures of ρ^wt^-ATPγS-Psu complexes. **a**, CryoEM reconstruction of ρ^wt^-ATPγS-Psu complex I. Six Psu dimers (violet and purple) bridge two open ρ^wt^ hexamers, ρ^wt^(A) and ρ^wt^(B). Individual subunits of ρ^wt^(A) and ρ^wt^(B) are colored in increasingly darker shades of gray from the center to the peripheries of the complex. **b**, Orthogonal views of ρ^wt^-ATPγS-Psu complex I. Psu dimers, cartoon representation; ρ^wt^, surface representation. Subunit coloring as in (**a**). ρ^wt^ subunits are designated by increasing Arabic numerals from the centers to the peripheries of the complex. **c**, CryoEM reconstruction of ρ^wt^-ATPγS-Psu complex II. Eight Psu dimers (violet and purple) bridge two open ρ^wt^ nonamers, ρ^wt^(A) and ρ^wt^(B). Portions of ρ^wt^(A) and ρ^wt^(B) equivalent to ρ^wt^-ATPγS-Psu complex I are shown in the same colors. ρ^wt^-ATPγS-Psu complex II can be envisaged to emerge from complex I by the additional of ρ^wt^ subunits and Psu dimers. Additional ρ^wt^ subunits added at the center of complex I, yellow; additional ρ^wt^ subunits added at the peripheries of complex I, light and dark green. **d**, Orthogonal views of ρ^wt^-ATPγS-Psu complex II. Psu dimers, cartoon representation; ρ^wt^, surface representation. Subunit coloring as in (**c**). The portion of complex II that is equivalent to complex I is shown in semi-transparent cartoon/surface representation, additional Psu dimers and ρ^wt^ subunits in complex II are shown in solid cartoon/surface representation. **e**, Scheme illustrating ρ^wt^-Psu interaction patterns in ρ^wt^-ATPγS-Psu complexes I and II. A dynamic ensemble of ρ^wt^-ATPγS-Psu complexes is supported by additional complexes observed in the cryoEM analysis (Extended Data Fig. 3).

**Fig. 2:**
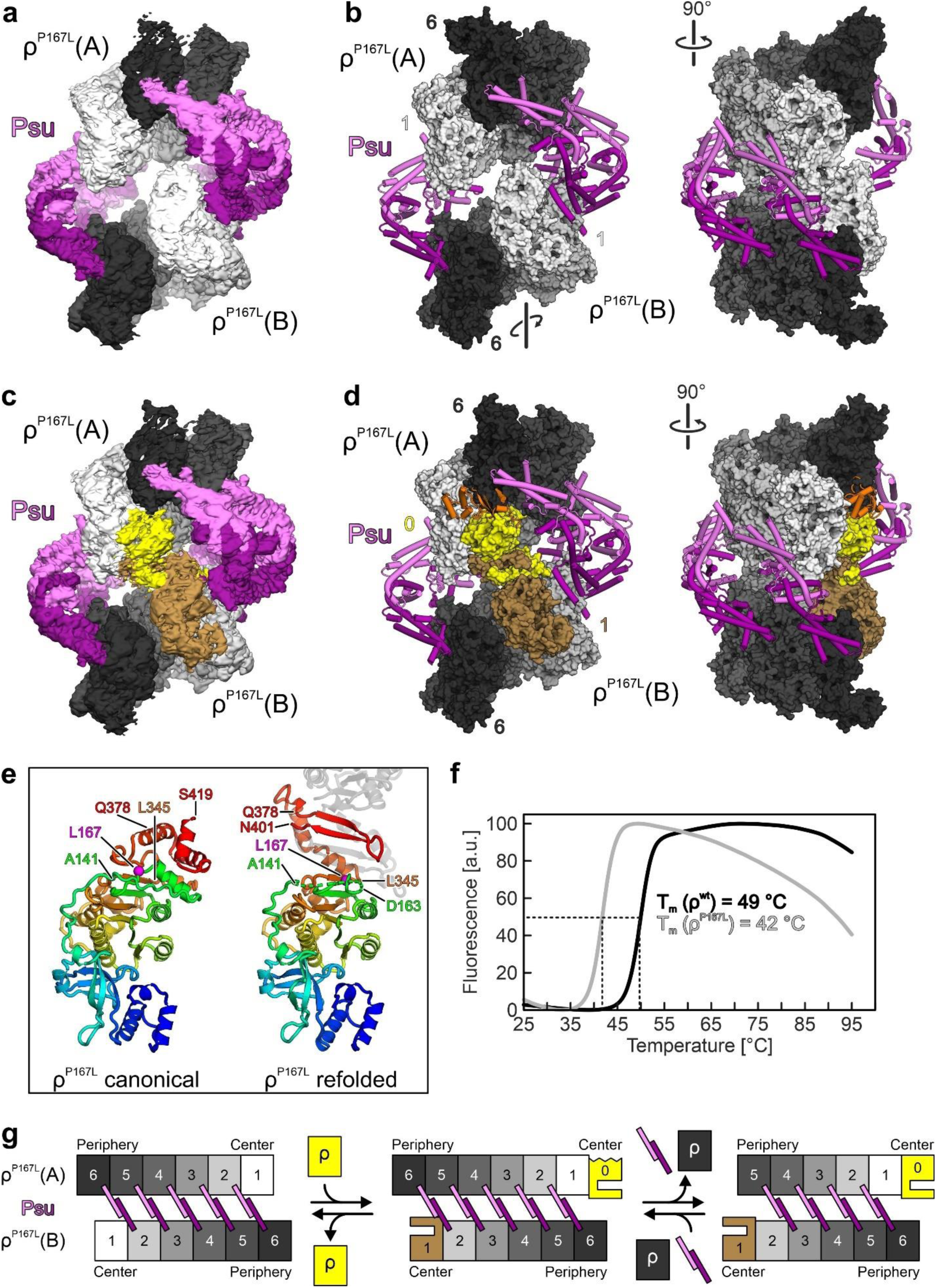
Structures of ρ^P167L^-ATPγS-Psu complexes. **a**, CryoEM reconstruction of ρ^P167L^-ATPγS-Psu complex I. Five Psu dimers (violet and purple) bridge two open ρ^P167L^ hexamers, ρ^P167L^(A) and ρ^P167L^(B). Subunit coloring as for ρ^wt^-ATPγS-Psu complex I (Fig. 1a). Rotation symbol, different rotational arrangement of ρ^P167L^(B) relative to ρ^P167L^(A), compared to ρ^wt^-ATPγS-Psu complex I. **b**, Orthogonal views of ρ^P167L^-ATPγS-Psu complex I. Psu dimers, cartoon representation; ρ^P167L^, surface representation. Subunit coloring as in (**a**). ρ^P167L^ subunits are designated by increasing Arabic numerals from the centers to the peripheries of the complex. **c**, CryoEM reconstruction of ρ^P167L^-ATPγS-Psu complex II. Compared to ρ^P167L^-ATPγS-Psu complex I, an additional ρ^P167L^ subunit (yellow) joined the complex, expanding ρ^P167L^(A) into a heptamer. The additional ρ^P167L^ subunit is refolded and engages in a domain-swapped interaction with ρ^P167L^ subunit 1 of the ρ^P167L^(B) hexamer (light brown). **d**, Orthogonal views of ρ^P167L^-ATPγS-Psu complex II. Psu dimers, cartoon representation; ρ^P167L^, surface representation. Subunit coloring as in (**c**). An N-terminal domain modeled in canonical conformation onto the additional ρ^P167L^ subunit (orange cartoon) would clash with the peripheral subunit of ρ^P167L^(A). **e**, Side-by-side comparison of a ρ^P167L^ subunit in canonical conformation (left) and in the refolded conformation (right). The models are colored blue to red from N- to C-terminus. For the refolded model, a second refolded ρ^P167L^ subunit, interacting *via* domain-swap in ρ^P167L^-ATPγS-Psu complex II, is shown as a gray, semi-transparent cartoon. Cα atom of L167, magenta sphere. Positions of landmark residues used to describe the rearrangement are indicated. **f**, Differential scanning fluorimetry monitoring fold stabilities of ρ^wt^ (black curve) and ρ^P167L^ (gray curve). T_m_ values were determined as the maxima of the first derivatives of the melting curves. Experiments were repeated independently at least three times with similar results. **g**, Scheme illustrating ρ^P167L^-Psu interaction patterns in ρ^P167L^-ATPγS-Psu complexes I (left) and II (center), and how an additional complex observed in the cryoEM analysis (right; Extended Data Fig. 6) could emerge from complex II.

**Fig. 3:**
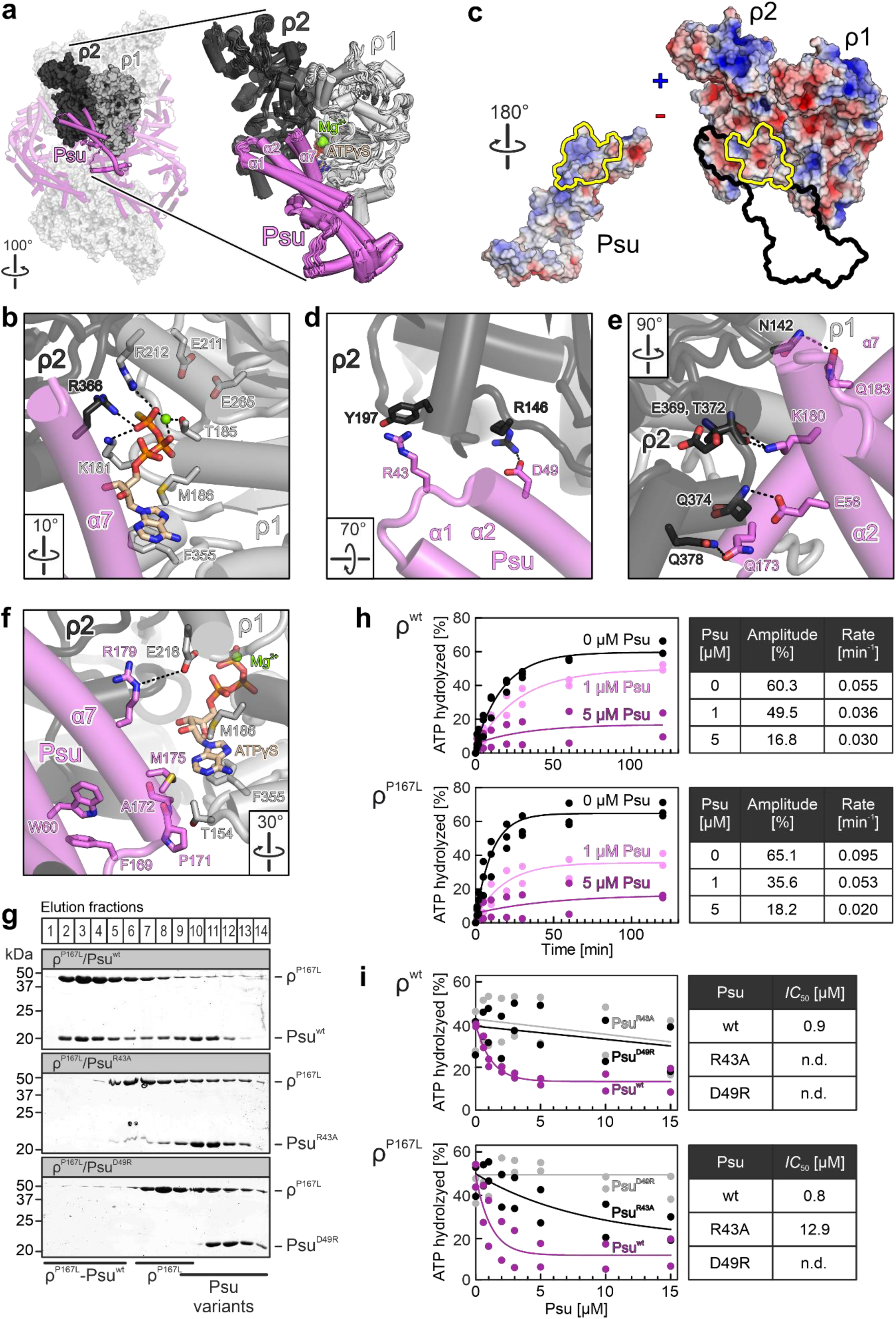
ρ-Psu interaction details. **a**, Left, ρ^wt^-ATPγS-Psu complex I with a building block comprising two neighboring ρ^wt^ subunits (ρ1 and ρ2) and an interacting Psu molecule highlighted in a solid representation. Right, overlay of all corresponding building blocks of the ρ^wt^-ATPγS-Psu complexes I and II. ρ-interacting helices of Psu are labeled. Bound ATPγS molecules and Mg^2+^ ions are shown as sticks and spheres, respectively, colored by atom type. In this and the following panels: ATPγS carbon, beige; nitrogen, blue; oxygen, red; sulfur, yellow; phosphorus, orange; magnesium, green. Rotation symbol, view relative to Fig. 1b, left. **b**, Details of ATPγS binding at the building blocks shown in (**a**). Relevant protein residues are shown as sticks and colored by atom type. In this and the following panels: carbon, as the respective protein subunit; black dashed lines, hydrogen bonds or salt bridges; rotation symbols, view relative to (**a**). **c**, Book-view on the interacting surfaces (yellow outlines) of Psu (left) and of two neighboring ρ^wt^ subunits (right), with the electrostatic potential (blue, positive; red, negative) mapped onto the protein surfaces. Black outline, border of the bound Psu molecule. **d,e,f**, Details of the ρ-Psu interaction within the building blocks shown in (**a**). **g**, SDS-PAGE analyses of analytical SEC elution fractions monitoring the interaction of ρ^P167L^ with Psu variants. Elution fractions are indicated at the top. The same elution fractions were analyzed for each run and were aligned below each other. Proteins and protein mixtures analyzed are indicated above each gel. Molecular mass markers are indicated on the left. Protein bands are identified on the right. Fractions containing isolated proteins or complexes are identified at the bottom. Experiments were repeated independently at least two times with similar results. **h**, Time traces monitoring ρ^wt^ (top) and ρ^P167L^ (bottom) RNA-stimulated ATPase activities in the absence of Psu or in the presence of 1 or 5 µM of Psu, recorded *via* TLC. Data were recorded as biological replicates, (ρ^wt^ or ρ^P167L^ alone, n = 3; ρ^wt^ or ρ^P167L^ plus Psu, n = 2). Data were fitted to a single-exponential equation; *A* = *A*_0_+(*A*_max_-*A*_0_)*(1-exp(-*k*\*t)); *A*, fraction of ATP hydrolyzed at time t; *A*_0_, fraction of ATP hydrolyzed at time zero; *A*_max_, fraction of ATP hydrolyzed at infinite time (amplitude); *k*, rate constant. Quantified amplitudes and rate constants are listed on the right. **i**, Inhibition of ρ^wt^ (top) and ρ^P167L^ (bottom) RNA-stimulated ATPase activities by increasing concentrations of Psu^wt^, Psu^R43A^ or Psu^D49R^. Data represent biological replicates, n = 2. Data were fitted to an [inhibitor] *vs.* response function, *A* = *A_min_*+(*A_max_*-*A_min_*)/(1+([inhibitor]/*IC_50_*)); *A*, fraction of ATP hydrolyzed at a given Psu (inhibitor) concentration; *A_min_* and *A_max_*, fitted minimum and maximum fractions of ATP hydrolyzed. Quantified *IC*_50_ values are listed on the right; n.d., not determined.

In both assemblies, two open ρ^wt^ rings, referred to as ρ^wt^(A) and ρ^wt^(B), face each other with their CTDs, while the ρ^wt^ NTDs are facing outwards (Fig. 1a-d). Multiple Psu dimers laterally bridge the two ρ^wt^ rings. The lower-oligomeric complex (complex I; global resolution 3.65 Å) contains two open ρ^wt^ hexamers and six Psu dimers (Fig. 1a,b). Designating individual subunits of ρ^wt^(A)/ρ^wt^(B) by increasing Arabic numerals from the inner-most to the outer-most subunits, Psu dimers bridge ρ^wt^(A) subunit 6 with ρ^wt^(B) subunit 1, ρ^wt^(A) subunit 5 with ρ^wt^(B) subunit 2, etc. (Fig. 1b). The highest-oligomeric complex (complex II; global resolution 4.25 Å) can be envisaged to emerge from complex I by the addition of one centrally positioned ρ^wt^ subunit and two peripheral ρ^wt^ subunits to each of the ρ^wt^(A) and ρ^wt^(B) hexamers of complex I (Fig. 1c-e). The expansion of the ρ^wt^(A)/ρ^wt^(B) complexes generates two additional pairs of ρ^wt^ subunits that can be bridged by two additional Psu dimers (Fig. 1d). The most peripheral ρ^wt^ subunits of the ρ^wt^(A)/ρ^wt^(B) rings in complex II are not stabilized by contacts to a Psu dimer, as no appropriately-spaced, unoccupied ρ^wt^ subunit remains in the opposite ρ^wt^ ring. These peripheral subunits are less well defined in the the cryoEM reconstructions, suggesting that they are more flexibly or sub-stoichiometrically connected (Fig. 1c). Due to the additional ρ^wt^ subunits incorporated at the center of complex II, the minimum spacing of the inner-most ρ^wt^(A)/ρ^wt^(B) subunits is decreased from ∼50 Å in complex I to ∼10 Å in complex II. As a consequence, the contacting Psu subunits also reciprocally interact *via* their C-terminal α7 helices (Fig. 1d). Several other complexes observed upon multi-particle 3D refinement represent intermediate oligomeric states (e.g., marked by dashed boxes in Extended Data Fig. 3), supporting the picture of a dynamic ensemble of ρ^wt^-ATPγS-Psu complexes in solution (Fig. 1e).

In isolated, open ρ hexamers (PDB ID: 1PV4)^32^, the average inter-subunit rise (8.6 Å) yields an overall ρ1-ρ6 rise (42.9 Å), which is too small to allow oligomerization beyond six ρ protomers. The ρ^wt^ hexamers in ρ^wt^-ATPγS-Psu complex I exhibit slightly increased inter-subunit and ρ^wt^1-ρ^wt^6 rise values (9.5 Å and 46.5 Å, respectively). The incorporation of additional ρ subunits supported by additional, bridging Psu dimers in complex II is facilitated by a further increase in the inter-subunit and ρ^wt^1-ρ^wt^6 rise values (12.0 Å and 57.1 Å, respectively). As a consequence, the helical pitch in complex II is large enough to allow expansion by additional ρ^wt^ protomers, which can associate at both ends of the central ρ^wt^ hexamers.

### Increased stability of a ρ^P167L^-Psu interaction is based on local refolding of the ρ^P167L^ variant

A proline at the 167-equivalent position of *E. coli* ρ is invariant across the bacterial kingdom (Extended Data Fig. 4a). A ρ variant harboring a P167L substitution has been observed to bind Psu more tightly than ρ^wt^.^33, 34^ Consistently, we observed ρ^P167L^-Psu complexes in analytical SEC even in the absence of nucleotides, and the more pronounced shift in the elution volume compared to ρ^wt^-nucleotide-Psu complexes suggested the formation of more stable complexes compared to ρ^wt^ (Extended Data Fig. 1b). However, in ρ^wt^-ATPγS-Psu structures, P167 of ρ does not engage in direct contacts to Psu that could explain these effects. To explore the structural basis underlying the higher stability of ρ^P167L^-Psu complexes, we also determined cryoEM structures of ρ^P167L^-ATPγS-Psu complexes.

**Fig. 4:**
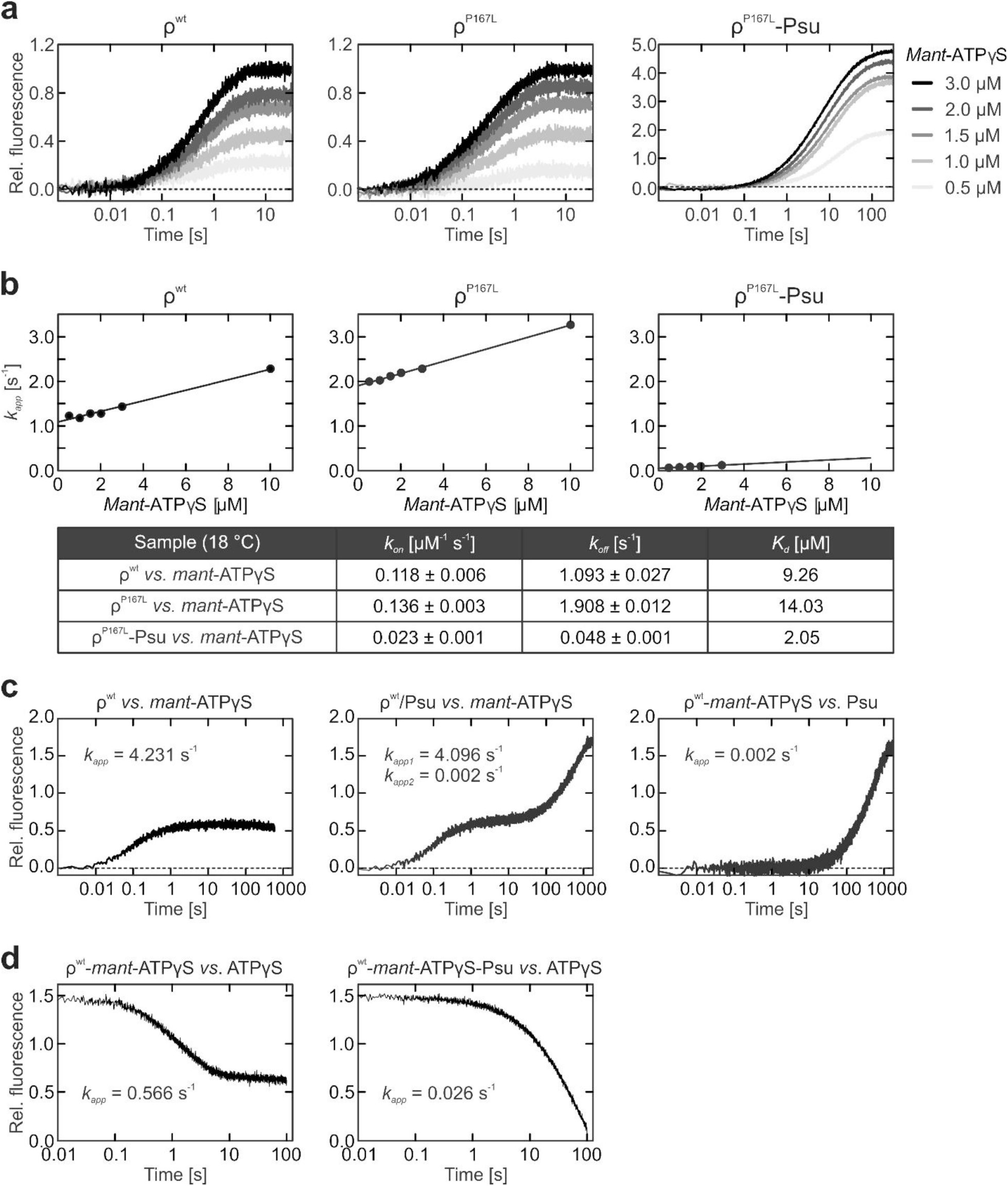
Effects of Psu on nucleotide binding by ρ. **a**, Time traces monitoring ρ^wt^ (left), ρ^P167L^ (middle) or ρ^P167L^-Psu (right) binding to increasing concentrations of *mant*-ATPγS (0.5 to 3 µM; colored from light grey to black), recorded *via* stopped-flow/FRET measurements. Data represent means, n = 3-6. Data were fit to a single exponential equation; *A* = *A_0_*+(*A_max_*-*A_0_*)*(1-exp(-*k_app_*\**t*)); *A*, fluorescence at time *t*; *A_max_*, final signal; *A_0_*, initial fluorescence signal; *k_app_*, characteristic time constant. **b**, Linear regression (*k_app_* = *k*_1_[*mant*-ATPγS] + *k*_−1_) of the apparent rate constants, *k*_app_, at increasing *mant*-ATPγS concentrations, derived by the single exponential fitting of the time-traces depicted in (**a**). The association (*k_on_*) and dissociation (*k_off_*) rate constants were derived from the slopes and y-intercepts. *K_d_*‘s were calculated as the ratios of the *k_off_* and *k_on_* values. **c**, Time traces monitoring ρ^wt^ (left) or ρ^wt^/Psu (middle) binding to *mant*-ATPγS, and of ρ^wt^-*mant*-ATPγS binding to Psu (right), recorded *via* stopped-flow/FRET measurements. Data represent means, n = 3-6. *k*_app_ values depicted in the graphs were derived by fitting of the data to a single exponential equation as in (**a**) (ρ^wt^ or ρ^wt^/Psu binding to *mant*-ATPγS), or by fitting of the data to a double exponential equation (ρ^wt^/Psu binding to *mant*-ATPγS); *A* = *A_0_*+*A_1_*(1-exp(-*k_app1_*\*t))+*A_2_*(1-exp(-*k_app2_*\*t); *A*, fluorescence at time *t*; *A_0_,* initial fluorescence signal; *A_1_* and *A_2_*, amplitudes of the signal change; *k_app1_* and *k_app2_*, characteristic time constants. **d**, Time traces monitoring the dissociation (replacement with ATPγS) of *mant*-ATPγS from ρ^wt^-*mant*-ATPγS-Psu (left) or ρ^P167L^-*mant*-ATPγS-Psu (right), recorded *via* stopped-flow/FRET measurements. Data represent means, n = 3-6. *k*_app_ values depicted in the graphs were derived by fitting of the data to a single-exponential equation; *A* = *A_min_*+(*A_0_*-*A_min_*)exp(-*k_app_*\**t*); *A*, fluorescence at time *t*; *A_min_*, final signal; *A_0_*, initial fluorescence signal; *k_app_*, characteristic time constant.

We obtained two main cryoEM reconstructions from ∼764,000 particle images (Extended Data Fig. 5 and 6; Supplementary Table 1). The first reconstruction (P167L-complex I; global resolution 3.09 Å) represents two open hexameric ρ^P167L^ rings, similar to ρ^wt^-ATPγS-Psu complex I, but with the ρ^P167L^(B) hexamer rotated by one subunit position relative to the ρ^P167L^(A) hexamer (Fig. 2a,b). As a consequence, the two opposed open ρ^P167L^ rings present only five pairs of ρ^P167L^ subunits that can be bridged by Psu dimers, which connect ρ^P167L^(A) subunit 6 with ρ^P167L^(B) subunit 2, ρ^P167L^(A) subunit 5 with ρ^P167L^(B) subunit 3, etc. (Fig. 2b). Assemblies with altered rotational orientation of Psu-bridged ρ^wt^ rings, and thus differing numbers of bridging Psu dimers, are also observed in the ensemble of the ρ^wt^-ATPγS-Psu complexes (e.g., marked by gray boxes in Extended Data Fig. 3).

**Fig. 5:**
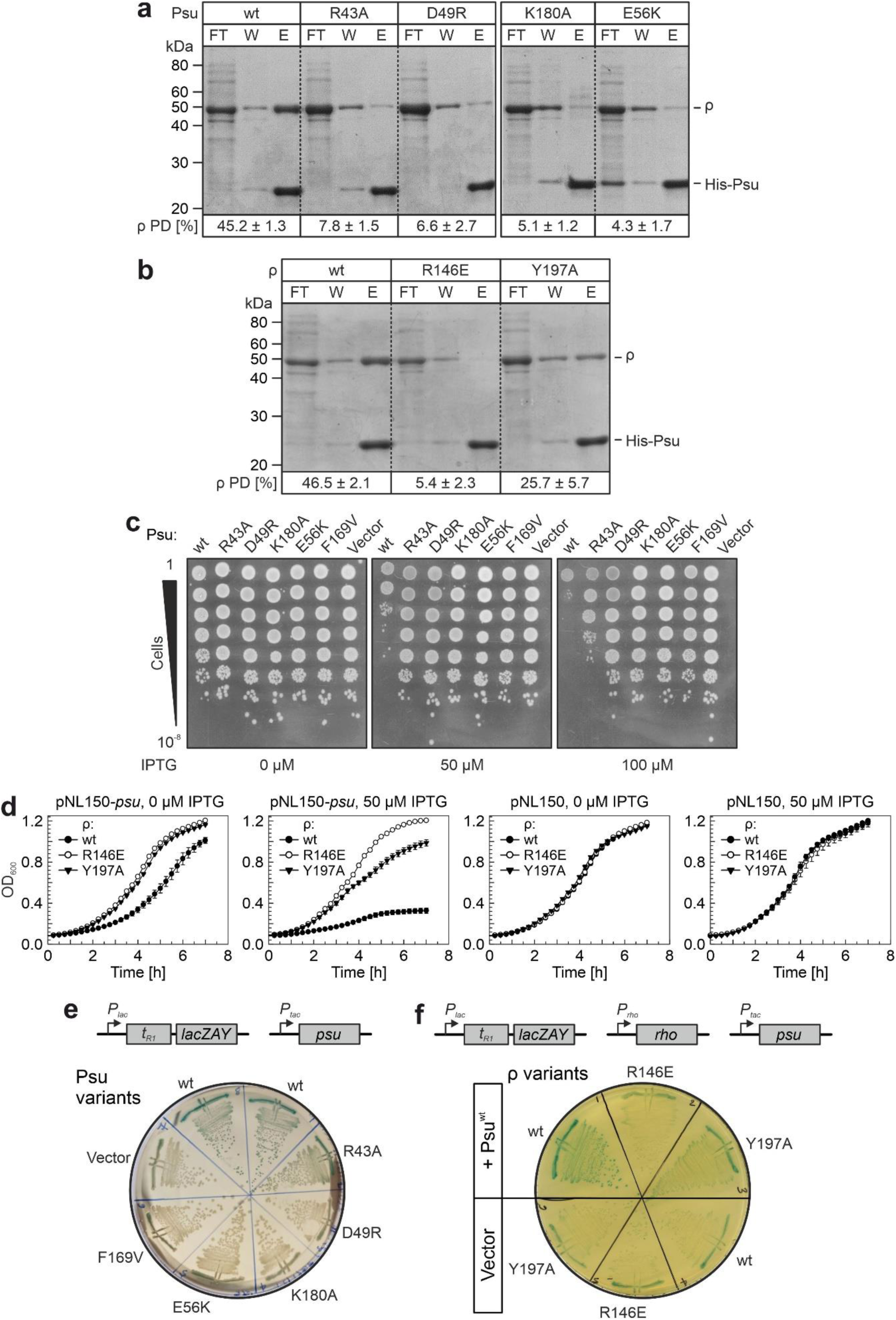
*In vivo* defects of Psu and ρ variants. **a**, *In vivo* binding defects of Psu variants. *E. coli* cell lysates after production of N-terminally His_6_-tagged Psu variants and ρ^wt^ were passed through Ni^2+^-NTA beads, and the flow-through (FT), wash (W), and eluted (E) fractions were collected. The fraction of pulled-down (PD) ρ^wt^ was calculated as (E/(FT+W+E))x100. Psu variants are indicated above the gels. Molecular mass markers are indicated on the left. Protein bands are identified on the right. Quantified data (ρ PD [%]) represent means ± SD; n = 3. **b**, *In vivo* binding defects of ρ variants. *E. coli* cell lysates after production of N-terminally His_6_-tagged Psu^wt^ and ρ variants were passed through Ni^2+^-NTA beads, and the flow-through (FT), wash (W), and eluted (E) fractions were collected. The fraction of pulled-down (PD) ρ variants was calculated as (E/(FT+W+E))x100. ρ variants are indicated above the gel. Molecular mass markers are indicated on the left. Protein bands are identified on the right. Quantified data (ρ PD [%]) represent means ± SD; n = 3. **c**, Serial dilutions of *E. coli* overnight cultures producing Psu variants (indicated above the plate images) were spotted on LB agar plates with different IPTG concentrations (indicated below the plate images). Serial dilutions are indicated on the left. Experiments were repeated independently at least two times with similar results. **d**, *E. coli* MG1655*ΔrhoΔrac* producing the indicated ρ variants were transformed with a plasmid guiding production of Psu^wt^ (first and second panel). As a control, strains were transformed with an empty vector (third and fourth panel). Growth curves were recorded in the absence of IPTG or in the presence of 50 µM IPTG, as indicated above the panels. Data points represent means ± SEM; n = 4-5. **e**, *E. coli* MG1655*ΔracΔlac* with a *P_lac_-t_R1_-lacZYA* reporter were transformed with plasmids guiding the production of the indicated Psu variants, or with an empty vector. Transformants were streaked on LB-X-gal agar plates supplemented with 15 µM IPTG. The reporter system produced blue-green colonies when the *in vivo* ρ-dependent termination function was inhibited. Experiments were repeated independently at least two times with similar results. **f**, *E. coli* MG1655*ΔrhoΔracΔlac* with a *P_lac_-t_R1_-lacZYA* reporter and producing the indicated ρ variants were further transformed with a plasmid guiding the production of Psu^wt^, or with an empty vector. Transformants were streaked on LB-X-gal agar plates supplemented with 15 µM IPTG. The reporter system produced blue-green colonies when the *in vivo* ρ-dependent termination function was inhibited. Experiments were repeated independently at least two times with similar results.

**Fig. 6:**
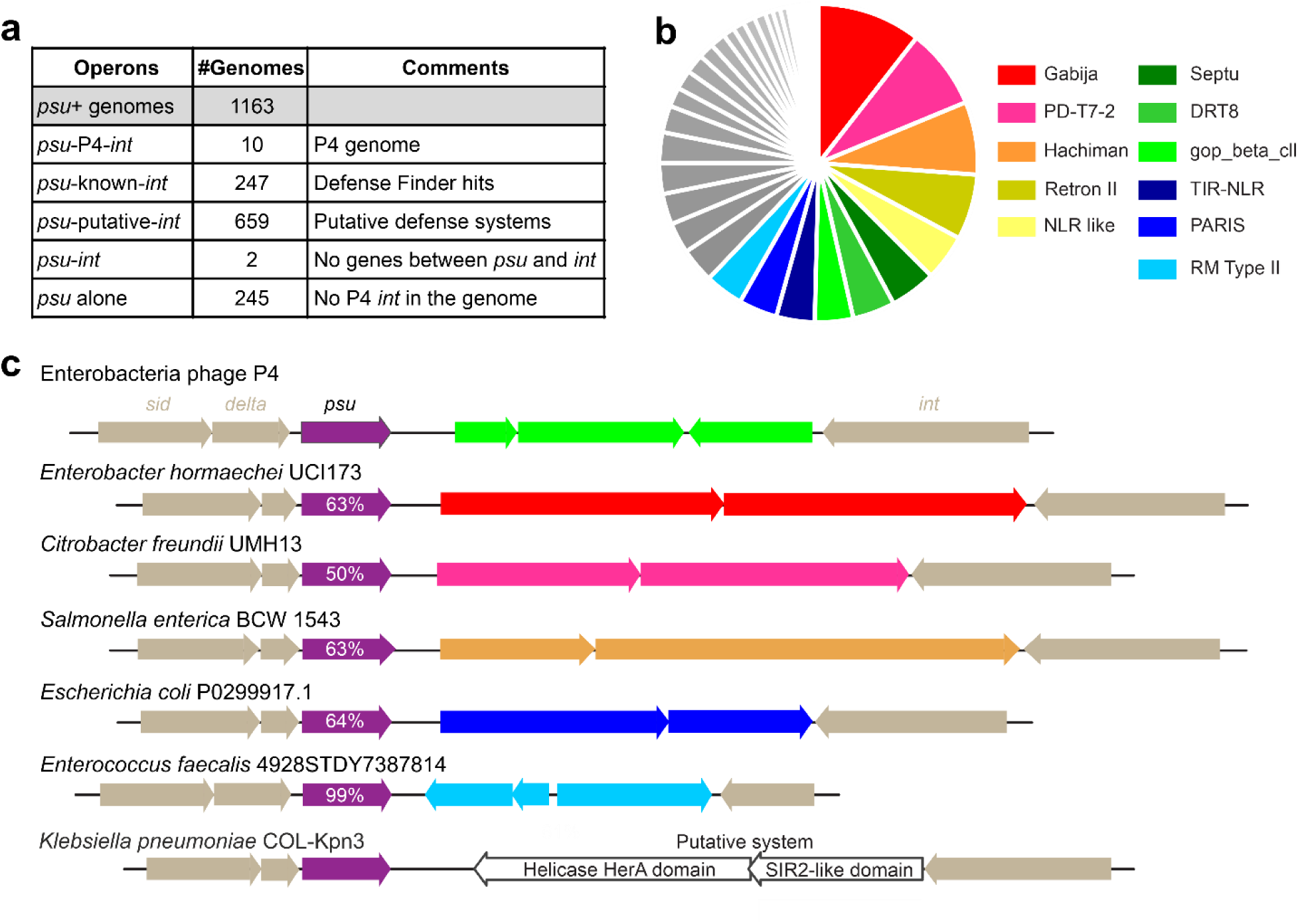
Psu as a marker of anti-phage defense systems. **a**, Defense system analysis of *psu*^+^ genomes. See Supplementary Dataset 1 for genomic context details. We note that the absence of the *int* gene could be due to incomplete genome assembly in some cases. **b**, The 37 classes of *psu-*associated defense systems identified by DefenseFinder are shown in a pie chart, with the eleven most abundant defense systems indicated in color; see Extended Data Fig. 8 for a complete list. **c**, Examples of *psu-*associated defense system loci; % identity to P4 Psu is shown. The known defense genes are colored as in **b**; a putative defense system with a SIR2-like domain is shown in white.

The second refined ρ^P167L^-ATPγS-Psu assembly (P167L-complex II; global resolution 2.95 Å) resembles the first arrangement but with an additional ρ^P167L^ subunit appended to the center of the ρ^P167L^(A) ring (Fig. 2c,d). Upon joining of this additional subunit, the helical pitch of the ρ^P167L^(A) ring does not increase, such that the NTD of the additional ρ^P167L^ subunit in its canonical conformation would clash with the most peripheral subunit of the ρ^P167L^(A) ring (Fig. 2d). As a consequence, the NTD of the additional ρ^P167L^ subunit is displaced and not defined in the cryoEM reconstruction. The CTD of the additional subunit in its canonical conformation would clash with the CTD of ρ^P167L^(B) subunit 1 (Fig. 2d); nevertheless, the additional subunit can be accommodated as both CTDs refold and form domain-swapped β-sheet arrangements (Fig. 2d,e). Upon refolding, the Psu interaction interface is altered, preventing Psu association with the central, refolded ρ^P167L^ subunits. Consequently, no additional Psu dimers can join as observed for the highest-oligomeric ρ^wt^-ATPγS-Psu complex II, preventing higher-level oligomerization of the ρ^P167L^ rings in this constellation.

In the refolded ρ^P167L^ protomers, the region preceding L167 (residues 141-163), encompassing the C-terminal part of the hinge region, becomes disordered, as indicated by a lack of interpretable cryoEM density (Fig. 2e). C-terminally of L167, the ρ^wt^ fold is maintained up to residue L345. In the following region up to residue Q378, helical elements are repositioned and changed in register. Between residues Q378 and N401, the helical elements of the canonical fold adopt an extended β-hairpin structure that constitutes the main swapped element (Fig. 2e). The swapped β-hairpins connect the central, six-stranded β-sheets of the CTDs of the domain-swapped ρ^P167L^ subunits into a continuous, 16-stranded sheet (Fig. 2e). Residues beyond N401 again lack interpretable cryoEM density.

Mutation-induced domain swapping has been described previously and is frequently associated with destabilization of the fold of the affected protein variants.^35^ Indeed, differential scanning fluorimetry revealed that the apparent melting temperature of ρ^P167L^ is decreased by about 7 °C compared to ρ^wt^ (Fig. 2f). Thus, the P167L substitution leads to a decrease in the fold stability of ρ subunits that allows refolding of part of the CTD. As a consequence, a more stable interlocking of ρ^P167L^ complexes is possible *via* the observed domain swapping, leading to the formation of overall more stable ρ^P167L^-Psu complexes.

Similar to ρ^wt^-ATPγS-Psu complexes, additional ρ^P167L^-ATPγS-Psu assemblies imaged can be considered to emerge from P167L-complexes I or II by the addition or dissociation of ρ^P167L^ subunits and Psu dimers. E.g., one such additional assembly contains two hexameric ρ^P167L^ rings interlocked by refolded, domain-swapped ρ^P167L^ subunits in the center and bridged by four Psu dimers (Fig. 2g, right); it could emerge from P167L-complex II upon dissociation of the peripheral ρ^P167L^(A)6 subunit and loss of the associated Psu dimer (marked by a dashed box in Extended Data Fig. 6). Due to the loss of the peripheral ρ^P167L^(A)6 subunit, the NTDs of both domain swapped central ρ^P167L^ protomers exhibit density for NTDs in the canonical conformation in this assembly. We suggest that ρ^P167L^ prefers to form Psu-bridged complexes that can be stabilized by domain swapping of the central, refolded ρ^P167L^ protomers.

### Psu binds across ρ nucleotide-binding pockets

ATPγS is bound similarly between all neighboring ρ CTDs in the ρ^wt^-ATPγS-Psu or ρ^P167L^-ATPγS-Psu assemblies (Fig. 3a). The nucleotides are bound as previously observed for open ρ complexes^36^ (Fig. 3b). Thus, the adenine base is sandwiched between F355 and M186, K181 contacts the γ-phosphate and T185 together with the β/γ-phosphates coordinates the Mg^2+^ ion (Fig. 3b). The γ-phosphate is additionally bound by R212 (arginine valve), and by R366 (arginine finger) of the adjacent ρ CTD (Fig. 3b). Compared to the hydrolytic situation, the catalytic glutamate (E211) and the Walker B aspartate (D265) are displaced from the ATPγS phosphates and there is no evidence for a bound attacking water molecule.

Except at the inner-most ρ(A/B) subunits of ρ^wt^-ATPγS-Psu complexes, which lack a preceding ρ subunit, each Psu molecule binds in a similar fashion between two neighboring ρ CTDs in all ρ^wt/P167L^-ATPγS-Psu assemblies (root-mean-square deviations of 0.6-2.0 Å for ∼8,100 atom pairs upon global, pairwise superposition of ρ-ρ-Psu sub-complexes; Fig. 3a). Psu interacts predominantly with one ρ CTD *via* the tip of its long coiled-coil (helices α1 and α2), the N-terminal part of helix α2 and the C-terminal helix α7. Psu helix α7 (residues 170-186) is placed in a cleft between the two neighboring ρ CTDs, covering the corresponding nucleotide-binding pocket (Fig. 3a). The interaction is supported by shape and electrostatic complementarity (Fig. 3c).

Psu R43 at the tip of the coiled-coil engages in cation-π interactions with Y197 of the main interacting ρ subunit (Fig. 3d). Psu D49 at the N-terminus of helix α2 (residues 45-91) forms an electrostatic interaction with ρ R146 (Fig. 3d). Psu E56 (helix α2) hydrogen bonds with the ρ Q374 backbone amide (Fig. 3e). Psu Q173, K180 and Q183 (helix α7) form hydrogen bonds with the ρ Q378 side chain, the backbone carbonyls of ρ E369/T372, and the side chain of ρ N142, respectively (Fig. 3e). Furthermore, there is a weak electrostatic interaction between Psu R179 (helix α7) and E218 of the adjacent ρ subunit (Fig. 3f). Finally, Psu P171, A172 and M175 (helix α7) engage in contacts with the T154-loop of the neighboring ρ subunit and cover the Watson-Crick side of the ATPγS base bound at that subunit, stabilizing it between ρ M186 and F355 (Fig. 3f). In the largest ρ^wt^-ATPγS-Psu complex II, additional contacts ensue between the dimerization region of the central Psu dimers and the NTDs of the central ρ subunits (Fig. 1d). The lack of conservation in the Psu-interacting residues in ρ across all bacteria (Extended Data Fig. 4a; Supplementary Dataset 1A) must be one of the reasons why the phage P4 host range is restricted to *Enterobacteriacea;* ∼99 % of *psu* genes are found in *Enterobacteriaceae* (Supplementary Dataset 1B). The conservation pattern of ρ-interacting residues in Psu supports the functional importance of helix α7-ρ contacts revealed by the structure: Psu residues P171, A172 and Q173 are highly conserved, and only K or R is found at position 180 (Extended Data Fig. 4b).

### ρ-contacting residues of Psu are required for complex formation and inhibition of ρ ATPase activity

Effects of previously investigated Psu and ρ variants^16, 19, 33, 37, 38^ are fully reconciled by our structures (Supplementary Table 2). To further test the importance of specific contacts observed in our structures, we exchanged Psu residues R43 and D49 (contacts to ρ Y197 and R146, respectively; Fig. 3d), to alanine and arginine, respectively, and tested the binding of these variants to ρ^P167L^ that forms complexes with Psu^wt^ independent of added nucleotides. Psu^R43A^ showed reduced binding to ρ^P167L^ in analytical SEC, as indicated by a reduced shift in elution volume compared to ρ^P167L^-Psu^wt^, while binding of Psu^D49R^ was completely abolished (Fig. 3g).

We next investigated Psu effects on ρ ATPase activity^38^ by measuring ATP hydrolysis by ρ^wt^ and ρ^P167L^, stimulated by RNA containing the *rut* site of the strong *λt_R1_* terminator. Reaction mixtures contained a fixed amount of ρ variants and increasing amounts of Psu^wt^ under pseudo-first-order conditions. Reactions were started by the addition of ATP, samples were taken at various time points, and nucleotides were separated by thin-layer chromatography (TLC; Source Data) and quantified. The time traces of hydrolyzed ATP could be fitted to a single exponential equation (Fig. 3h). The final percentage of hydrolyzed ATP (reaction amplitude) was comparable for ρ^wt^ and ρ^P167L^, but the ATP turnover rate of ρ^P167L^ was approximately doubled compared to ρ^wt^ (Fig. 3h). For both ρ variants, addition of increasing amounts of Psu^wt^ (Psu^wt^ dimer:ρ hexamer 10:1 and 50:1) led to the stepwise decrease of the ρ ATPase rate (1.5-fold and 2-fold, respectively, for ρ^wt^; 2-fold and 5-fold, respectively, for ρ^P167L^) and amplitude (by 20 % and 70 %, respectively, for ρ^wt^; by 55 % and 70 %, respectively, for ρ^P167L^; Fig. 3h). Psu^wt^ inhibited RNA-stimulated ATPase activities of ρ^wt^ and ρ^P167L^ with comparable *IC_50_* values (0.9 µM and 0.8 µM, respectively; Fig. 3i). Psu^R43A^ was partially defective, with an *IC_50_* value 10-fold higher (12.9 µM) than for Psu^wt^ (Fig. 3i), whereas Psu^D49R^ was fully defective in inhibiting the ρ^P167L^ ATPase, and neither Psu variant significantly affected the ATPase activity of ρ^wt^ (Fig. 3i). Together, the above results confirm the importance of specific Psu residues in establishing contacts to ρ that inhibit the RNA-stimulated ATPase activity of ρ, fully consistent with our structures.

### Psu effectively inhibits nucleotide trafficking on ρ

Previously, a reduced ATP affinity of ρ was observed in the presence of Psu based on a cross linking assay.^38^ As Psu helix α7 covers nucleotide-binding sites in our ρ^wt/P167L^-ATPγS-Psu structures (Fig. 3a,b), Psu may hinder nucleotide binding to and release from ρ. To examine this possibility, we monitored *mant*-ATPγS binding *via* Förster resonance energy transfer (FRET) from ρ W381 in the vicinity of the nucleotide-binding pockets in a stopped-flow device. To this end, we quickly mixed constant amounts of ρ variants or pre-assembled ρ^P167L^-Psu complex with increasing concentrations of *mant*-ATPγS at 18 °C. For each *mant*-ATPγS concentration, the apparent rate constants (*k_app_*) of binding were determined by fitting the time traces of fluorescence changes to a single exponential equation (Fig. 4a). The *k_app_* values were plotted against the nucleotide concentrations and fitted to a linear regression function to estimate the association rate constant (*k_on_*; slope) and the dissociation rate constant (*k_off_*, y-intercept; Fig. 4b). Dissociation constants (*K_d_*) were calculated as the ratios of the *k_off_* and *k_on_* values (Fig. 4b).

In the absence of Psu, ρ^wt^ and ρ^P167L^ exhibited similar *K_d_*‘s for *mant*-ATPγS (9.3 µM and 14.0 µM, respectively; Fig. 4b). The slightly lower *mant*-ATPγS affinity of ρ^P167L^ is due to a two-fold higher dissociation rate (Fig. 4b), which likely arises because the residue substitution renders the nucleotide-binding site more dynamic, as indicated by the lower thermal stability of ρ^P167L^ (Fig. 2f). The increased nucleotide dissociation constant may enable faster ATP turnover, consistent with the observed elevated ATPase rate of ρ^P167L^ compared to ρ^wt^ (Fig. 3h).

The time-dependent change in the FRET signal observed with ρ^P167L^-Psu was almost five-fold stronger than for ρ^P167L^ alone (Fig. 4b). We attribute this effect to bound Psu, which contributes additional FRET donors (W20, W60) in the vicinity of the ρ^P167L^ nucleotide-binding pockets, close to or below the tryptophan-*mant* Förster radius of ∼25 Å^39^, thus enhancing the energy transfer to the *mant*-moiety. In cryoEM reconstructions of ρ^wt/P167L^-ATPγS-Psu complexes, we observed density features suggesting covalent attachment of ATPγS to C117 of Psu upon prolonged incubation. In principle, a similar covalent attachment of *mant*-ATPγS could contribute to the observed increase in FRET signal, as Psu residues W20, W60 and W133 reside within a distance of ∼25 Å or less of C117. However, mixing of Psu and *mant*-ATPγS did not lead to a significant FRET signal in the timeframe of the experiment.

The ρ^P167L^-Psu complex exhibited a ∼seven-fold higher *mant*-ATPγS affinity compared to ρ^P167L^ alone (*K_d_* = 2.1 µM; Fig. 4b). The *mant*-ATPγS association and dissociation rates of the complex were more than five-fold and ∼40-fold lower, respectively, compared to ρ^P167L^ alone (Fig. 4b). The drastic decrease of the *mant*-ATPγS dissociation rate in the presence of Psu suggests that, once nucleotide is bound, Psu effectively hinders its release.

Consistent with our observations that ρ^wt^ does not form stable complexes with Psu in the absence of nucleotides (Extended Data Fig. 1a), we did not observe Psu-dependent changes in the FRET signal when quickly mixing *mant*-ATPγS with ρ^wt^/Psu at 18 °C. We, therefore, increased the reaction temperature to 37 °C to accelerate binding events and acquired longer time traces to monitor possible slower interactions. Under these conditions, we observed an expected temperature-dependent increase of the *k_app_* for *mant*-ATPγS binding (about four-fold compared to 18 °C; Fig. 4c, first panel). In the presence of Psu, two phases could be clearly distinguished and the time-dependent FRET signal could be fitted to a double-exponential function (Fig. 4c, second panel). The apparent rate of the fast phase (*k_app1_*) was equivalent to that of *mant*-ATPγS binding to ρ^wt^ alone, indicating *mant*-ATPγS binding to unmodified ρ^wt^ (Fig. 4c, first and second panels). As the slow phase (*k_app2_*) was associated with a further increase in the FRET signal, similar to what was observed with ρ^P167L^ in the presence of Psu, it likely monitored the subsequent binding of Psu to the ρ^wt^-*mant*-ATPγS complex. To test this notion, we pre-assembled ρ^wt^-*mant*-ATPγS complex and quickly mixed it with Psu in the stopped-flow instrument. The resulting time dependent FRET signal could be fitted with a single exponential equation, which indicated an apparent rate of Psu binding in perfect correspondence to the slow phase (*k_app2_*) of the ρ^wt^/Psu *vs. mant*-ATPγS measurement (Fig. 4c, second and third panels). Thus, the *k_app_* values derived from the individual ρ^wt^ to *mant*-ATPγS and ρ^wt^-*mant*-ATPγS to Psu binding assays are equivalent to the fast rate constant, *k_app1_*, and the slow rate constant, *k_app2_*, respectively, derived from the double exponential fit of the ρ^wt^/Psu to *mant*-ATPγS binding assay. Finally, we mixed an excess of non-labeled ATPγS with pre-formed ρ^wt^-*mant*-ATPγS or ρ^wt^-*mant*-ATPγS-Psu complexes to quantify the *mant*-ATPγS dissociation rates. Presence of Psu decreased the apparent dissociation rate constant more than 20-fold (Fig. 4d).

Collectively, these results confirm the higher affinity of ρ^P167L^ to Psu compared to ρ^wt^; corroborate that Psu binding to ρ^wt^, but not to ρ^P167L^, is dependent upon prior nucleotide binding to ρ^wt^; and support the notion that Psu helix α7 obstructs access to, and exit from, the ρ nucleotide binding pockets. The reduced nucleotide binding and release in the presence of Psu likely contributes to the observed Psu-mediated inhibition of the ρ ATPase.

### Psu does not affect RNA binding at PBSes but inhibits SBS binding of ρ^P167L^

Stable RNA binding at the ρ SBS and RNA-stimulated ATPase activity of ρ require ring closure.^22^ In our ρ^wt/P167L^-ATPγS-Psu complex structures, ρ invariably adopts an open ring conformation, and we showed that Psu inhibits ρ ATPase (Fig. 3), suggesting that Psu may also interfere with RNA-SBS interactions. In contrast, ρ NTDs are invariably turned outwards in the structures, in principle granting RNAs access to the ρ PBSes. In agreement with the latter notion, Psu did not affect ρ binding to the PBS ligand poly(dC).^38^

To test potential effects of Psu on ρ-nucleic acid interactions in more detail, we used fluorescence anisotropy assays.^40^ ρ PBSes bind pyrimidine-rich DNA or RNA while the ρ SBS exclusively engages RNA.^41^ Thus, for probing PBS binding, we used a fluorescein-labeled DNA oligomer, dC15-FAM. For testing RNA binding to the SBS, we used a fluorescein-labeled RNA oligomer, rU12-FAM, after saturating the PBSes with unlabeled dC15. ADP-BeF3 was included in the assays to stabilize ρ-Psu interactions. PBS and SBS affinities to the respective nucleic acid probes were very similar for ρ^wt^ and ρ^P167L^ (Extended Data Fig. 7a-d). Increasing concentrations of Psu did not affect dC15-FAM binding at the PBSes of either ρ variant (Extended Data Fig. 7a,b), consistent with our cryoEM structures revealing unobstructed PBSes in the Psu-bridged ρ complexes. However, the prior addition of Psu to ρ^P167L^-dC_15_ inhibited rU_12_-FAM binding at the SBS (Extended Data Fig. 7d; *IC_50_* = 2.8 µM), while Psu failed to displace rU_12_-FAM from pre formed ρ^P167L^-dC_15_-rU_12_-FAM complexes (Extended Data Fig. 7e). In contrast, Psu inhibited the ρ^wt^ SBS binding only very weakly (Extended Data Fig. 7c; *IC_50_* = 299 µM). Together with the effect of Psu on RNA-stimulated ρ ATPase, these observations suggest that while Psu stabilizes the open conformation of ρ^wt^ or ρ^P167L^, it still allows binding of RNA at the center of the open ρ^wt^ complexes, but inhibits RNA binding to the center of open ρ^P167L^ complexes. Possibly, the refolded central ρ^P167L^ subunits observed in ρ^P167L^-ATPγS-Psu complexes limit access of RNA to the centers of the corresponding Psu-bound ρ^P167L^ complexes.

### Psu variants defective in ρ binding exhibit anti-termination defects *in vivo*

To test the importance of interactions observed in our ρ^wt/P167L^-ATPγS-Psu complex structures in cells, we monitored the effects of selected ρ and Psu variants under *in vivo* conditions. Apart from interaction-deficient Psu^R43A^ and Psu^D49R^ (Fig. 3d,g), we constructed Psu^K180A^, in which hydrogen bonds of K180 with E369 and T372 of ρ are abolished (Fig. 3e). We also generated ρ^Y197A^, in which the cation-π stacking of Y197 with R43 of Psu is not possible (Fig. 3d). Selected, previously investigated Psu and ρ variants^33, 37, 38^ served as controls: Psu^E56K^, which cannot sustain an interaction with the backbone amide of Q374 (Fig. 3e); Psu^F169V^, in which the proper positioning of helix α7 is likely compromised; and ρ^R146E^, in which a salt bridge to Psu D49 is destroyed (Fig. 3d).

To monitor potential interaction defects of ρ or Psu variants *in vivo*, we co-produced ρ variants and N-terminally His_6_-tagged Psu variants in the *E. coli* BL21(DE3) strain, prepared cell lysates of mid-log phase cultures, and conducted pull-down assays using Ni^2+^-NTA beads, as described previously^33, 38^. About 45 % of ρ^wt^ was pulled down *via* Psu^wt^, while all tested Psu variants strongly reduced co-precipitation (< 8%; Fig. 5a). Thus, all Psu variants expected to be defective in ρ binding based on our structures indeed failed to bind ρ^wt^ efficiently *in vivo*. Likewise, only about 5 % or about 25 % of ρ^R146E^ or ρ^Y197A^, respectively, were co-precipitated by Psu^wt^ (Fig. 5b), consistent with ρ-Psu contacts observed in our structures.

Over-production of Psu^wt^ or of Psu variants that bind and inhibit ρ leads to severe growth and ρ-dependent termination defects.^38, 42^ We, therefore, produced Psu^wt^ and variants from an IPTG inducible *P*_tac_ promoter in *E. coli*, and spotted serial dilutions of saturated cultures in the absence or presence of 50 µM or 100 µM IPTG. Production of Psu^wt^ caused severe growth defects, whereas the growth of most strains that produced Psu variants defective in ρ binding was unaffected (Fig. 5c). Only Psu^R43A^ production led to mild growth defects in the presence of 100 µM IPTG (Fig. 5c), consistent with only partial disruption of the ρ^P167L^ interaction in analytical SEC (Fig. 3g) and partial inhibition of ρ ATPase activity by this variant *in vitro* (Fig. 3i)

Next, we co-transformed the MG1655 strain that harbors a chromosomal deletion of the *rho* gene with the *Ptac*-Psu^wt^ plasmid and plasmids producing ρ variants. Upon induction of Psu^wt^ by 50 µM IPTG, strains producing ρ^R146E^ or ρ^Y197A^ showed no or reduced growth defects, respectively (Fig. 5d). These results are fully in line with the lost or partially reduced interactions of Psu^wt^ with ρ^R146E^ or ρ^Y197A^ suggested by our structures and observed in pull-down assays (Fig. 5b).

We then investigated effects on ρ-dependent termination *in vivo* using a reporter system in which the *lacZ* gene was positioned downstream of a strong ρ-dependent terminator, λ*t_R1_* (Fig. 5e). As β-galactosidase production from the reporter depends on the suppression of ρ-dependent termination, colonies will appear white or pale blue on LB-X-gal plates if ρ is functional, and will appear blue-green if ρ function is inhibited. When Psu^wt^ was expressed in the reporter strain, colonies appeared blue-green, indicating inhibition of ρ. Consistent with partial defects in ρ binding observed *in vitro* (Fig. 3g) and in pull-down assays (Fig. 5a), Psu^R43A^-producing colonies were light blue, whereas production of all Psu variants defective in ρ binding led to white or pale blue colonies (Fig. 5e).

Finally, *E. coli* cells with the *lacZ* reporter were transformed with plasmids expressing ρ^wt^ or variants, and subsequently, the chromosomal *rho* gene was deleted. The resulting strains were then transformed with plasmids encoding Psu^wt^ (or empty plasmid as control), and the colonies were streaked on LB-X-gal plates. Colonies producing ρ^wt^ in the presence of Psu^wt^ appeared blue-green, indicating Psu^wt^-mediated inhibition of ρ^wt^-dependent termination (Fig. 5f). In contrast, colonies producing ρ^Y197A^ or ρ^R146E^, defective in Psu^wt^ binding based on our structures, appeared pale blue (Fig. 5f), consistent with these ρ variants being partially resistant to inhibition by Psu^wt^.

### P4-like mobile elements encoding Psu are hubs for bacterial defense genes

The role of Psu may not be limited to P4 infection. Recent studies revealed that P4-like mobile elements are hotspots for bacterial defense systems.^43–45^ Almost half of all *E. coli* genomes were found to carry P4-like elements adjacent to anti-phage systems that included abortive infection (Abi), restriction-modification (RM) and other defense modules.^44^ In P4, Psu is encoded in the *sid*-*δ*-*psu* operon and, consistent with Psu suppressing polarity, activates expression of *δ* and *psu*, but not of *sid*.^18^ As a general anti-terminator, Psu may also upregulate associated defense clusters by conferring protection against premature termination by ρ. To explore this possibility, we investigated 1,163 bacterial genomes encoding Psu (Extended Data Fig. 8) and used DefenseFinder^46^ to detect putative defense systems in the *psu* neighborhood. This analysis revealed a known defense system inserted between the *psu* and *int* (integrase) genes of P4 in more than a quarter (247) of *psu*^+^ genomes (Fig. 6a), consistent with acquisition through horizontal transfer^44^. These defense systems are compact (∼3 kb), presumably reflecting the P4 genome size limitations, and astonishingly diverse, representing 37 known classes (Fig. 6b; Extended Data Fig. 8; Supplementary Dataset 1CD). Interestingly, one system, RM type II, is found in *Enterococcus faecalis*, a Firmicute (Fig. 6c); the latter example likely represents a very recent horizontal transfer event because of the near identity of the Psu protein to its P4 homolog.

The catalog of known defense systems is far from complete. Bioinformatic screens based on gene syntax identified many novel putative defense gene clusters, which have been shown to encode active defense elements^47–49^. We thus used analogous guilt-by-association criteria to search for putative defense systems in the *psu-int* region. This analysis revealed 659 inserts that were not identified by DefenseFinder (Fig. 6a; Supplementary Dataset 1E) but share the compact size of known defense systems (Fig. 6c). An HMM search against the Pfam database for the proteins encoded in these loci identified 767 hits from 107 Pfam families that contain nucleases, methylases, toxin-antitoxin and Abi systems, and other components commonly present in phage defense systems (Supplemental Dataset 1). This analysis suggests that a considerable fraction of the P4 “dark matter” is also involved in transferring defense genes across host genomes, and that new P4-linked defense systems can be found *in silico*, followed by experimental validation. For example, our analysis identified a sirtuin (Sir2)-HerA helicase module adjacent to Psu in *Klebsiella pneumoniae* (Fig. 6c). Although this module was not recognized by DefenseFinder, an analogous Sir2-HerA system from *Paenibacillus* sp. has been recently shown to trigger abortive infection and cell death^50^ *via* the Sir2-mediated depletion of the cellular pool of NAD^+^, a signature activity of Sir2 in different defense contexts^50–52^.

Many defense genes are codirectional with Psu (Fig. 6c), potentially extending the *psu* operon and falling under its cognate polarity control^18^. However, given that Psu works in *trans*, including on chromosomal genes, it may activate any gene that is subject to ρ-mediated termination, a common mechanism of xenogeneic silencing^8^.

## Discussion

P4 is a paradigmatic satellite phage that relies on its helper phage, P2, to produce functional virions. The P4-encoded Psu plays two vital roles in P4 infection: it stabilizes the Sid-hijacked P2 capsid proteins to selectively package the P4 DNA^53^, and it inhibits ρ-mediated silencing of phage genes^18^. Here, we elucidated the structural basis and molecular mechanisms underlying the Psu mediated inhibition of ρ. Our cryoEM structures revealed that Psu instigates hyper-oligomerization of ρ on two levels. First, multiple Psu dimers cross-strut two open ρ complexes; and second, lower-level ρ-Psu complexes can incorporate additional ρ subunits, expanding ρ to at least the nonamer stage, in part aided by the concomitant incorporation of additional Psu dimers.

During capsid morphogenesis, two P4 proteins, Psu and its distant paralog Sid, constrict the P2 capsid^53^; the Psu/Sid-modified capsid can only accommodate a much smaller P4 genome, one-third the length of P2 (11 kb *versus* 33 kb). In a high-resolution reconstruction of the P4 pro capsid, Sid dimers are seen bound across hexamers of gpN, the P2 capsid protein.^54^ A low resolution structure of the mature P4 capsid, in which Psu replaces Sid, suggested that the structurally similar Psu dimer likewise caps gpN hexamers.^55^ Guided by the Psu-gpN interaction as well as by mutational and cross-linking studies, a model of a ρ-Psu complex has been proposed, in which a Psu dimer also caps a closed ρ ring.^33^ Our cryoEM structures of ρ-ATPγS-Psu complexes revealed a decisively different mode of ρ-Psu complex formation, fully in line with previous (Supplementary Table 2) and present mutational analyses of the ρ-Psu interaction, the estimated ∼1:1 subunit stoichiometry in ρ-Psu complexes^33, 42^, and additional biochemical data presented here. First, Psu blocks the nucleotide-binding pockets on ρ – we show that Psu reduces the binding and release of nucleotides to and from ρ (Fig. 4). Second, ρ adopts an open, inactive conformation in all complex structures with Psu – we demonstrate that Psu efficiently inhibits ρ ATPase activity (Fig. 3). Third, in all our structures the NTDs of the two Psu-bridged ρ complexes are facing outwards – we show that Psu does not inhibit nucleic acid binding to the ρ PBSes (Extended Data Fig. 7a). Finally, our functional analyses addressing the effects of Psu and ρ variants on ρ-Psu interaction in cells, cell viability, and ρ-mediated termination *in vivo* (Fig. 5) are fully consistent with our cryoEM structures.

Psu-mediated inhibition of ρ ring closure would inhibit the ρ translocase activity, and thus ρ-mediated termination *via* the tethered tracking pathway^25^. Open and expanded ρ complexes bridged by Psu would likely also be incompatible with ρ transferring NusA/NusG-modified ECs into a pre-termination state, as recently imaged^27, 28^. While ρ-Psu complexes might still engage NusA/NusG-ECs, Psu would most likely hinder the conformational changes needed to inactivate RNAP or inhibit late stages of termination/hybrid unwinding, when ρ is expected to engage and translocate on RNA ^27^. We posit that the Psu-mediated blockade of the nucleotide-binding pockets of ρ contributes to the inhibition of the ρ ATPase activity. Interestingly, ATPγS (and thus presumably ATP) supports Psu binding to ρ (Supplemenraty Fig. 1). We attribute this effect of ATPγS (ATP) to the stabilization of the Psu contact surface on the ρ CTDs and direct contacts of Psu to residues surrounding the nucleotide-binding pockets (Fig. 3f). Once bound, however, Psu inhibits the ρ ATPase activity. Thus, Psu exhibits typical features of an uncompetitive inhibitor of the ρ ATPase.

ρ activity is regulated on multiple levels. On the one hand, interactions with nucleotides, RNAs, or NusG shift the equilibrium between open and closed hexameric states and thereby regulate ρ ATPase and RNA translocase activities.^22, 56^ On the other hand, RNAP-associated NusA^57, 58^, an RNAP-trailing ribosome^59–61^, RNAP-bound elements of nascent RNA^14, 15^, RNAP-bound specialized transcription factors^11–13^, or multi-factorial RNAP-bound RNA-protein complexes^9, 10^ can delay or prevent the attack of ECs by ρ. Furthermore, the RNA chaperone Hfq can counteract ρ by depositing ρ-inhibitory sRNAs on nascent transcripts.^4, 62^ Results presented here underscore an additional mechanism *via* which ρ can be regulated, *i.e.*, factor-dependent modulation of its oligomerization state. This mechanism capitalizes on the intrinsic oligomerization dynamics of ρ, with the hexamer representing the only state in which ρ can terminate transcription. Depending on the concentration and conditions, ρ can dissociate into non-functional, lower oligomeric states, down to the monomer level, or form dodecamers, albeit exhibiting a different configuration, as seen in complexes with Psu.^63–65^ While some proteins, such as *Vibrio cholerae* YaeO/Rof, may stabilize ρ in a hypo-oligomerized state^66^, Psu stabilizes hyper-oligomerized, inactive states. Apart from interacting proteins, some ρ orthologs have acquired additional regions or domains that facilitate ρ regulation *via* modulation of its oligomeric state: *Clostridium botulinum* ρ has an additional, N-terminal prion-like domain that promotes the formation of amyloid-like aggregates^67^, while an intrinsically disordered region in *Bacteroides thetaiotaomicron* ρ can promote phase separation associated with increased termination activity^68^. We posit that condition-dependent activation or inactivation of ρ through changes in its oligomeric or aggregation state may be a widespread regulatory strategy.

Modulation of the oligomerization state as a regulatory mechanism has been observed in other molecular systems, including in eukaryotes, and is of growing interest among drug developers.^69^. For example, plant or mammalian ASPL/PUX1 proteins can disassemble the essential and multi functional AAA+ ATPase CDC48/p97^70–72^, whereas biparatopic designed ankyrin repeat proteins can induce apoptosis by oligomerization of human epidermal growth factor receptor-2 in nanoscopic membrane domains^73^. The development of oligomerization-modulating compounds can be fueled by structural knowledge of the target protein in different oligomeric states dependent on a cellular interaction partner. Inhibitory peptides (shiftides) of HIV-1 integrase, for instance, have been designed based on the interacting lens epithelium-derived growth factor/p75, which shift HIV-1 integrase into an inactive, tetrameric state.^74^ Results presented here could thus inspire the development of novel ρ-inhibitory substances. Peptides (and eventually peptide-derived compounds) designed based on ρ-contacting regions of Psu might not only block the nucleotide binding pockets on ρ but may also facilitate ρ hyper-oligomerization. While short peptides or low molecular-weight compounds will not support the dimerization of two open ρ rings akin to Psu, substances that bind to and increase the pitch of open ρ hexamers may conceivably allow expansion into inactive ρ filaments. Notably, Psu-derived, ρ-inhibiting peptides that are active *in vivo* have already been designed^75^, but the structural basis of their modes of action remains to be explored.

Our findings also suggest that Psu may play a hitherto unknown role in antiviral defense. Phage-defense systems encoded on mobile elements are pan-genomic “guns for hire”^76^ that bacteria deploy when under attack by phages^77^. In rapid exchange through horizontal transfer, these elements shape bacterial adaptation and evolution, and can inform the development of new phage therapies. P4-like elements are integrated into about 30 % of enterobacterial genomes with a high probability for retained functionality, and many of these elements carry phage-defense systems.^43–45^ Several of these systems have been shown to protect *E. coli* from lytic phages, including λ, P1, and T7, while not restricting P2^44^, implying the existence of mutualistic relationships between parasitic satellites and their helpers. Our *in silico* analysis suggests that Psu proteins may serve as built-in anti-terminators that insulate the associated defense genes from ρ-mediated silencing. Curiously, the RM systems, which dominate the bacterial defense arsenal,^46^ are underrepresented among Psu-associated defense genes (Fig. 6b), which are instead enriched in Abi modules, such as Gabija^78^, retrons^79^, and PD-T7-2^45^. In fact, Psu itself is a potent toxin that, if present at high levels, triggers cell death in diverse bacteria^80^.

While supplying the cell with antiviral weapons, a Psu/defense cassette also poses a threat to its very existence. The expression of Abi systems must be tightly controlled to ensure that a suicide program is triggered only when the first line of cellular defense has been breached.^81^ ρ limits the expression of the *psu* gene ^18^ and could be a key part of the silencing mechanism. Furthermore, ρ may have evolved to limit its sensitivity to Psu: P167L substitution stabilizes ρ-Psu interactions ^33, 34^, and we traced this enhanced stability to a tendency of ρ^P167L^ to refold and form domain-swapped dimers at the center of ρ^P167L^-Psu complexes. Interestingly, we find an invariant proline at the 167-equivalent position in bacterial ρ proteins (Extended Data Fig. 4a). Notably, ρ^P167L^ does not exhibit defects in ATPase activity or termination *in vitro* and supports *E.coli* growth^33^; thus, the invariance of P167 could hint at an evolutionary pressure to limit ρ-Psu affinity and thereby balance Psu-dependent expression of phage defense genes and cellular toxicity of Psu.

## Methods

### Recombinant protein production and purification

Primers used for molecular cloning are listed in Supplementary Table 3. Plasmids used for protein production or as templates for *in vitro* transcription are listed in Supplementary Table 4. Genes encoding Psu variants (R43A and D49R) were produced by the single-primer quick mutagenesis method^82^. For protein production, *E coli* BL21 RIL cells were transformed with corresponding plasmids. ρ^wt^, ρ^P167L^ and Psu^wt^ were produced and purified as previously described^27, 33, 83^. Briefly, lysates containing non-tagged ρ^wt^ or ρ^P167L^ were loaded on a heparin column equilibrated in 50 mM Tris-HCl, pH 7.5, 50 mM NaCl, 5 % (v/v) glycerol, 1 mM DTT. The proteins were eluted with a gradient to 1 M NaCl, and the fractions containing ρ^wt^ or ρ^P167L^ were pooled, concentrated and loaded on a Superdex 200 column for SEC in 10 mM Tris-HCl, pH 7.5, 150 mM NaCl, 1 mM DTT. Psu^wt^ or variants were precipitated with 25 % final concentration of ammonium sulfate for 1 h at RT, and pelleted by 15,000 x g centrifugation at 4 °C for 20 min. The pellet was resuspended in 20 mM Tris-HCl, pH 8.9, 150 mM NaCl, 5 % (v/v) glycerol, 0.1 mM EDTA, 1 mM DTT and passed through heparin and MonoQ chromatography columns to clear contaminants. The flow-through was subjected to MonoS cation-exchange chromatography, eluted with a gradient to 1 M NaCl and the fractions containing Psu variants were pooled, concentrated and loaded on a on a Superdex 75 column for SEC in 20 mM Tris-HCl, pH 8.0, 100 mM NaCl, 10 % (v/v) glycerol, 0.1 mM EDTA, 1 mM DTT.

### *Rut* RNA production

A DNA template for *in vitro* transcription, harboring a T7 RNAP promoter and the sequence encoding the λ*tR1 rut* site, was produced from plasmid pUC18-λ*t_R1_*-*rut* (Supplementary Table 4) by PCR with a reverse primer containing two 2’-O-methylated nucleotides at the 5’-terminus, for decreasing possible 3’-end run-off products.^84, 85^. T7 RNAP-based *in vitro* transcription was performed as described previously.^10^ The produced λ*tR1 rut* site RNA was treated with DNase I and purified by strong anion-exchange (MonoQ, GE Healthcare) and size-exclusion chromatography (Superdex 75, GE Healthcare) in 10 mM HEPES-NaOH pH 7.5, 50 mM NaCl. The purity of the RNA was assessed by urea-PAGE, stained with ethidium bromide.

### Cryogenic electron microscopy

ρ^wt^-ATPγS-Psu and ρ^P167L^-ATPγS-Psu complexes were freshly prepared in 10 mM TRIS-HCl, pH 7.5, 150 mM NaCl, 5 mM MgCl_2_ (ρ^wt^-ATPγS-Psu), or 10 mM TRIS-HCl, pH 8.0, 120 mM KOAc, 5 mM Mg(OAc)_2_, 5 µM ZnCl_2_ (ρ^P167L^-ATPγS-Psu), concentrated to 5.5 mg/ml using a 100 kDa ultra-centrifugal filter (Merck) and supplemented with 1 mM ATPγS. 3.8 µl of the samples were applied to glow-discharged holey carbon R1.2/1.3 copper grids (Quantifoil Microtools) and plunge frozen in liquid ethane using a Vitrobot Mark IV (Thermo Fisher) equilibrated at 10 °C and 100 % humidity. Micrographs were recorded on a FEI Titan Krios G3i transmission electron microscope operated at 300 kV with a Falcon 3EC detector. Movies were recorded for 40.57 s, accumulating a total electron flux of ∼40 el/Å^2^ in counting mode at a calibrated pixel size of 0.832 Å/px distributed over 33 fractions.

### CryoEM data analysis

All image analysis steps were done with cryoSPARC (version 3.2.2)^86^. Movie alignment was done with patch motion correction generating Fourier-cropped micrographs (pixel size 1.664 Å/px), and CTF estimation was conducted by Patch CTF. Class averages of manually selected particle images were used to generate an initial template for reference-based particle picking from 5,986/2,723 micrographs (ρ^wt^-ATPγS-Psu/ρ^P167L^-ATPγS-Psu). 1,815,462/823,078 particle images were extracted with a box size of 224 px and Fourier-cropped to 112 px for initial analysis. Reference-free 2D classification was used to select 747,325/763,878 particle images for further analysis. *Ab initio* reconstruction using a small subset of particles was conducted to generate an initial 3D reference for subsequent classification by 3D heterogeneous refinement or 3D variability analysis. Local motion correction was applied to extract particle images at full spatial resolution with a box size of 448 px followed by global and local CTF refinement. Due to considerable structural and compositional flexibility of the complexes, classification was not straightforward. Instead, similar structures appeared multiple times during classification, which were then combined for final homogeneous refinement. For the ρ^wt^-ATPγS-Psu sample, classes of 15,407 particle images (4.25 Å) and 73,056 particle images (3.65 Å) were finally reconstructed by non uniform (NU) refinement. For ρ^P167L^-ATPγS-Psu, 317,635 and 273,935 particle images were selected for final NU refinement, yielding reconstructions at global resolutions of 2.95 Å and 3.09 Å, respectively.

### Model building, refinement and analysis

Crystal structures of ρ (PDB ID: 1PV4)^32^ and Psu (PDB ID: 3RX6)^83^ were manually placed in the cryoEM reconstructions and each protomer was adjusted by rigid body fitting and segmental real-space refinement using Coot (version 0.9.6)^87, 88^. For higher-oligomeric complexes, additional ρ subunits and Psu dimers were added to unoccupied regions of the cryoEM reconstructions and adjusted as above. The models were refined by iterative rounds of real space refinement in PHENIX (version 1.20_4459)^89, 90^ and manual adjustment in Coot. The structural models were evaluated with MolProbity (version 4.5.1)^91^. Structure figures were prepared with PyMOL (version 2.4.0, Schrödinger) and ChimeraX^92^.

### Analytical size-exclusion chromatography

Analytical SEC-based interaction analyses were performed in 10 mM TRIS-HCl, pH 7.5, 150 mM NaCl, 5 mM MgCl_2_, 1 mM DTT. 300 pmol ρ^wt/P167L^ hexamer and 1200 pmol Psu dimer variants were mixed in a final reaction volume of 50 µl. After incubation of the mixtures at 32 °C for 10 min, the samples were loaded on a Superose 6 increase 3.2/300 analytical size exclusion column (Cytiva). 50 µl fractions were collected and analyzed by SDS-PAGE and visualized by Coomassie staining.

### Differential scanning fluorimetry

Differential scanning fluorimetry was conducted in a 96-well plate in a plate reader combined with a Mx3005P thermocycler (Stratgene). ρ variants were prepared at a final concentration of 2 µM in 10 mM Tris-HCl, pH 7.5, 150 mM NaCl, and 1 mM DTT, supplemented with 10x SYPRO orange in a final volume of 20 µl. The temperature was increased at a rate of 1 °C/min from 25 °C to 95 °C, and the fluorescence emission was monitored at each step. The fluorescence intensity was plotted as a function of temperature. The thermal melting temperature was determined from the melting profile as the temperature at which the fluorescence intensity reached 50 % of the highest fluorescence intensity.

### ATPase assays

TLC-based ATPase assays were performed using [α-^32^P]ATP (Hartmann Analytic). To quantify RNA-stimulated ATPase activity, 100 nM ρ variants, in isolation or supplemented with 1 µM or 5 µM Psu, were mixed with 20 µM *rut* RNA in 10 mM Tris-HCl pH 7.5, 100 mM NaCl, 5 mM MgCl_2_, 1 mM DTT. The mixtures were incubated with 1 mM ATP, supplemented with [α-³²P]ATP, at 37°C for up to 2 h. 2 µl of the samples were withdrawn at selected time points and the reactions were quenched with 6 µl of 100 mM EDTA, pH 8.0. 3 x 1 µl of samples were spotted on a PEI cellulose TLC membrane and chromatographed with 1 M acetic acid, 0.5 M LiCl, 20 % (v/v) ethanol. The corresponding ATP and ADP spots were visualized using a Storm 860 phosphorimager (GMI) and quantified using ImageQuant software (version 5.2; Cytiva). To test the effect of Psu variants on ρ ATPase activity, increasing concentrations of Psu variants, up to 15 µM, were mixed with 100 nM ρ variants, and subsequently with 20 µM *rut* RNA, in 10 mM Tris-HCl pH 7.5, 100 mM NaCl, 5 mM MgCl_2_, 1 mM DTT. The mixtures were incubated with 1 mM ATP, supplemented with [α-³²P]ATP, at 37 °C for 1 h. Subsequently, the experiments were conducted as above. Data from two biological replicates were plotted and analyzed using Prism software (version 9.0.2; GraphPad). ρ^wt^ or ρ^P167L^ ATPase activity was calculated as the percentage of hydrolyzed ATP over time, by fitting quantified data to the equation *A* = *A_0_*+(*A_max_*-*A_0_*)*(1-exp(-*k*\**t*)), in which *A_0_* is the fraction of ATP hydrolyzed at time zero; *A_max_* is the the fraction of ATP hydrolyzed at infinite time; and *k* is the rate constant. To quantify the effect of Psu on ρ^wt^ or ρ^P167L^ ATPase activity, data were fitted to an inhibitor-vs.-response function: *A* = *A_min_*+(*A_max_*-*A_min_*)/(1+([inhibitor]/IC*_50_*)), in which *A* is the fraction of ATP hydrolyzed at a given Psu (inhibitor) concentration; and *A_min_* and *A_max_* are the fitted minimum and maximum fractions of ATP hydrolyzed.

### Stopped-flow/FRET-based analysis of nucleotide binding

Nucleotide binding studies were conducted on an SX-20MV stopped-flow/fluorescence spectrometer (Applied Photophysics). Tryptophan residues near the ATP binding pockets served as FRET donors and were excited at 280 nm. A 420 nm cut-off filter was used for detecting the emission by the acceptor, *mant*-ATPγS. All experiments were performed with 100 nM ρ^wt^ or ρ^P167L^, 500 nM Psu and/or 2 µM *mant*-ATPγS in 10 mM Tris-HCl, pH 7.5, 150 mM NaCl, 5 mM MgCl*_2_*. For quantifying nucleotide association, data was acquired at 18 °C or 37 °C by fast mixing of equal volumes (60 µl) of the reactants (syringe 1, ρ^wt^ or ρ^P167L^ in isolation or in the presence of Psu; syringe 2, *mant*-ATPγS). FRET signals were recorded for up to 30 min. For detecting the binding of Psu to ρ^wt^ or ρ^P167L^ pre-bound to *mant*-ATPγS, the experiment was performed at 37 °C by rapidly mixing equal volumes (60 µl) of the reactants (syringe 1, ρ^wt^ or ρ^P167L^ and *mant*-ATPγS; syringe 2, Psu) and the monitoring fluorescence changes over 30 min. Data were acquired in logarithmic sampling mode and visualized using the Pro-Data Viewer software package (version 2.5; Applied Photophysics). The final curves were obtained by averaging 3 to 6 individual traces and analyzed in Prism software (version 9.0.2; GraphPad). Unless stated otherwise, data were fitted to a single exponential function: *A* = *A_0_*+(*A_max_*-*A_0_*)*(1-exp(-*k_app_*\**t*)); *A*, fluorescence at time *t*; *A_max_*, final signal; *A_0_*, initial fluorescence signal; *k_app_*, characteristic time constant. For ρ^wt^ binding to *mant*-ATPγS and the subsequent association of Psu, a double exponential function was used: *A* = *A_0_*+*A_1_*(1-exp(-*k_app1_*\*t))+*A_2_*(1-exp(-*k_app2_*\*t); *A*, fluorescence at time *t*; *A_0_,* initial fluorescence signal; *A_1_* and *A_2_*, amplitudes of the signal change; *k_app1_* and *k_app2_*, characteristic time constants. Dependencies of the apparent rate constants on nucleotide concentration were fitted by a linear equation, *k_app_* = *k_1_*[*mant*-ATPγS] + *k_−1_*; *k_1_*, nucleotide association rate constant (derived from the slope); *k_−1_*, nucleotide dissociation rate constant (derived from the Y-axis intercept). For *mant*-ATPγS dissociation, data were fitted to a single exponential function: *A* = *A_min_*+(*A_0_*-*A_min_*)exp(-*k_app_*\**t*); *A*, fluorescence at time *t*; *A_min_*, final signal; *A_0_*, initial fluorescence signal; *k_app_*, characteristic time constant.

### Nucleic acid binding assays

Nucleic acid binding to ρ^wt^ or ρ^P167L^ PBS or SBS was tested *via* fluorescence depolarization-based assays.^40^ For PBS binding, 5 µM FAM-labeled dC_15_ oligo (Eurofins) were mixed with increasing amounts of ρ^wt^ or ρ^P167L^ (0 to 8 µM final hexamer concentration) in 20 mM HEPES pH 7.5, 150 mM KCl, 5 % (v/v) glycerol, 5 mM MgCl_2_, 0.5 mM TCEP. For SBS binding, ρ^wt^ or ρ^P167L^ PBSes were first saturated with 10 µM non-labeled dC_15_. Increasing amounts of PBS-blocked ρ^wt^ or ρ^P167L^ (0 to 16 µM final hexamer concentration) were then mixed with 2 mM ADP-BeF_3_ and 2 µM FAM-labeled rU_12_ oligo (Eurofins). For examining the effect of Psu on nucleic acid binding to the ρ^wt^ or ρ^P167L^ PBS or SBS, 1 µM of ρ^wt^ or ρ^P167L^ hexamers were mixed with increasing amounts of Psu (0 to 30 µM final dimer concentration). Subsequently, experiments were conducted as above. The fluorescence anisotropy was recorded in OptiPlateTM 384-well plates (PerkinElmer) using a Spark Multimode Microplate reader (Tecan; excitation wavelength, 485 nm; detected emission wavelength, 530 nm). Two technical replicates were averaged for each sample and the data were analyzed with Prism software (version 9.0.2; GraphPad). To quantify ρ^wt^ or ρ^P167L^ PBS or SBS binding, data were fitted to a single exponential Hill function; *A* = *A_max_*[protein]^h^/(*K* ^h^ + [protein]^h^); in which *A_max_* is the fitted maximum of nucleic acid bound; *K_d_* is the dissociation constant; and h is the Hill coefficient. For quantification of the effects of Psu on ρ^wt^ or ρ^P167L^ PBS or SBS binding, data were fitted to an inhibitor *vs.* response function: *A* = *A_min_*+(*A_max_*-*A_min_*)/(1+([inhibitor]/IC*_50_*)); in which *A* is the anisotropy signal at a given concentration of Psu (inhibitor); and *A_min_* and *A_max_* are the fitted minimum and maximum of nucleic acid bound.

### Generation of *rho* and *psu* mutants for *in vivo* assays

A low-copy-number pCL1920 plasmid guiding the production of ρ^Y197A^ was constructed by site directed mutagenesis of a pCL1920 plasmid encoding ρ^wt^ (pRS649)^93^. pNL150 plasmids guiding the production of Psu^R43A^, Psu^D49R^ or Psu^K180A^ were constructed by site-directed mutagenesis of a pNL150 plasmid encoding Psu^wt^ (pRS1117)^37^. Mutations were confirmed by sequencing.

### Bacterial growth assays

To monitor possible growth defects of *rho* mutants in the absence and presence of Psu, *E. coli* RS1309 (MG1655*ΔrhoΔrac*) strain was transformed with pCL1920 plasmids expressing wt or mutant *rho* and were plated on LB in the absence of IPTG. After removal of the shelter plasmid expressing wt *rho*, the strains were transformed with empty pNL150 or with pNL150 expressing wt *psu*. The transformed colonies were streaked on LB agar plates supplemented with 0 µM or 50 µM IPTG and incubated at 37 °C. Serial dilutions of the overnight cultures of each transformed strain were spotted onto LB agar plates supplemented with 0, 50 or 100 µM IPTG.

To monitor possible growth defects of the cells expressing *psu* mutants, *E. coli* MG1655 (RS1263) strain was transformed with pNL150 plasmids expressing either wt or mutant *psu*. The transformed colonies were streaked on LB agar plates supplemented with 0 µM or 50 µM IPTG.

To record growth curves, overnight cultures of the transformed strains were cultured in 96-well microtiter plates in the absence of IPTG or in the presence of 50 µM IPTG. The growth was monitored by measuring the absorbance at 660 nm using a Spectramax M5 plate reader. The curves were plotted using SigmaPlot software (version 13).

### Pull-down assays

The pET28b (*kan^R^*) plasmids guiding the production of Psu variants with an N-terminal His_6_ tag, and pET21b (*amp^R^*) plasmids guiding the production of ρ variants were co-transformed in *E. coli* BL21(DE3) cells. The transformed colonies were used to inoculate 5 ml of LB medium, and cultures were grown at 37 °C for ∼3 h. The 5 ml cultures were then added to 100 ml LB medium and further incubated until the OD_600_ reached ∼0.3. The cultures were then induced with 0.1 mM IPTG for 3 h. Cells were then lysed in 100 mM NaH_2_PO_4_, pH 7.0, 100 mM NaCl,10 mM imidazole, 1 mg/ml lysozyme, 10 mg/ml PMSF. The lysates were passed through Ni^2+^-NTA columns (Qiagen), washed with 100 mM NaH_2_PO_4_, pH 7.0, 100 mM NaCl, 20 mM imidazole, and the bound protein was eluted with 100 mM NaH_2_PO_4_, pH 7.0, 100 mM NaCl, 500 mM imidazole. Identical volumes of all buffers were used in each step. Identical volumes of the flow-through (FT), wash (W) and eluted fractions (E) were separated *via* SDS-PAGE. The amounts of ρ and Psu proteins in each of the FT, W and E fractions were quantified by ImageQuant (version 5.2; Cytiva) of the corresponding bands of the Coomassie-stained gels.

### *In vivo* ρ-dependent transcription termination assays

A *lacZ* reporter system was used for studying *in vivo* ρ-dependent transcription termination. *E. coli* RS2047 strain, containing the reporter cassette, *P_lac_-t_R1_-lacZYA*, in the form of a λRS45 lysogen, was transformed with pCL1920 plasmids that guide the production of ρ variants. Subsequently, cells were transduced with *rho::kan^R^* lysate to remove the chromosomal copy of *rho*. Finally, the modified strains were transformed with an empty pNL150 vector or with pNL150 expressing *psu*. Transformed colonies were streaked on LB-X-gal agar plates supplemented with 0 or 5 µM IPTG, and were incubated at 37 °C for 14 to 16 h.

To study the effect of *psu* mutants on ρ-dependent transcription termination *in vivo*, the RS2047 strain was transformed with empty pNL150 or with pNL150 expressing wt or mutant *psu*. The transformed colonies were streaked on LB-X-gal agar plates supplemented with 0 or 15 µM IPTG and were incubated at 37 °C for 12 to 14 hrs.

### ρ conservation analysis

To analyze the conservation of ρ, a representative database was compiled (Supplementary Dataset 1A). The GTDB^94^ bacterial taxonomy list (Released April 08, 2022) was downloaded. Genomes labeled as gtdb_type_species_of_genus and gtdb_representative were kept. Then the genome assembly level was inspected. Only complete genomes were used to build the database. The new dataset covers 41 phyla including 649 genus. The protein sequences of all genomes were downloaded from NCBI. To identify ρ, Pfam model Rho_RNA_bind (PF07497) was searched against the representative database using hmmsearch (v. 3.3)^95^ with a bit score of 27. The identified ρ-like proteins were further confirmed by searching ρ NCBI HMM model (NF006886.1) against them using hmmsearch with cutoff 315. Finally, 632 ρ sequences are collected.

The length of ρ sequences ranges from 355 to 857 residues. Many of them have long insertions. To detect local homologies, multiple sequence alignment (MSA) was done by Dialign2^96^ with the default settings. The MSA was indexed according to *E. coli* MG1655 ρ (NP_418230.1). The sequence conservation score was calculated using Protein Residue Conservation Prediction^97^ with property entropy option. Sequence logo was generated using WebLogo (v. 3.7.8)^98^.

### Compiling Psu sequences

The Pfam model (PF07455) of Psu was searched against Reference Proteomes (v.2021_04) using HmmerWeb (v. 2.41.2)^99^ with bit scores 21.2 and E-values 10^-4^. However, only 28 sequences were found, one from Enterobacteria phage P4 and 27 from Enterobacterales. To obtain a better picture of Psu diversity, we launched a ‘greedy search’: 1) searching Psu Pfam model against uniprotKB (v.2021_04) using HmmerWeb^99^ with bit score 21.2 and e-value 10^-4^; 2) Jackhmmer searching with Enterobacteria phage P4 Psu reference sequence (NCBI ID: NP_042044.1) as a query against uniprotKB (v.2021_04) using HmmerWeb^99^. A total of seven Iterations were done; 3) blastp searching with Psu reference sequence as a query against uniprotKB (v.2022_04) using UniProt website^100^ with e-value 10^-4^; 4) blastp searching Psu reference sequence against Reference proteins using NCBI website^101^ with e-value 10^-4^. Max target sequences option was set to 5000; 5) blastp searching Psu reference sequence against Non-redundant protein sequences using NCBI website^101^ with e-value 10^-4^. The search was limited to the Virus superkingdom (Taxonomy ID: 10239). Max target sequences option was set to 5000.

All sequences from the ‘greedy search’ were downloaded from NCBI and pooled together. An all-vs-all blastp was performed using BLAST+ (v. 2.9.0)^102^ with e-value 10^-9^. A Psu similarity network was built based on e-value and visualized in Cytoscape (v. 3.9.1)^103^. Sequences from two Psu-like groups in the network (Extended Data Fig. 8) were fed into Batch CD-Search^104^ to confirm the presence of the Psu superfamily (NCBI ID: cl06476). Sequences with > 80% coverage of the Psu superfamily and sequences with coverage < 80% but having no ’incomplete’ flags were selected. Finally, 2625 Psu sequences were kept (Supplementary Dataset 1B). NCBI taxonomy ID was first assigned to the Psu sequences using accession2taxid file (https://ftp.ncbi.nlm.nih.gov/pub/taxonomy/accession2taxid/). Then NCBI taxonomy was replaced with GTDB taxonomy^94^. As the reference sequences are not organism-specific, the taxonomy was assigned to the genus level.

### Psu conservation

The compiled Psu sequences from Pseudomonadota were clustered at 90 % (-c 0.9 option) using cd-hit (v. 4.8.1)^105^. Then two sequences from Firmicutes and three sequences from the virus were added into the pool. A total of 124 sequences were aligned using MUSCLE (v. 5.1) with default settings (https://doi.org/10.1101/2021.06.20.449169). The multiple sequence alignment of Psu was indexed according to Psu reference sequence before generating sequence logo using WebLogo (v. 3.7.8)^98^.

### Identification of anti-phage defense systems

The accession numbers of ten neighboring genes on both sides of *psu* were fetched by TREND^106^ and the sequences were downloaded from NCBI. A total of 1,163 *psu*-containing genomes were investigated. The presence of Enterobacteria phage P4-like genes among the neighboring genes was determined using blastp. The proteins from phage P4 reference genome (NCBI ID: GCF_000846325.1) were used as queries, and e-value threshold was set to 10^-5^. To find known anti-phage defense system, all neighboring proteins were submitted to DefenseFinder^46^. The results were checked manually. Gene clusters located between *psu* and *int* genes that were not identified by DefenseFinder were considered putative defense systems (Supplementary Dataset 1E) and genes from putative defense systems were annotated (Supplementary Dataset 1G,H) by searching against Pfam-A database (v. 35.0) using hmmscan (v. 3.2.1)^95^.

## Data availability

CryoEM reconstructions have been deposited in the Electron Microscopy Data Bank (https://www.ebi.ac.uk/pdbe/emdb) under accession codes EMD-17637 (https://www.ebi.ac.uk/pdbe/entry/emdb/EMD-17637; ρ^wt^-ATPγS-Psu complex I), EMD-17639 (https://www.ebi.ac.uk/pdbe/entry/emdb/EMD-17639; ρ^wt^-ATPγS-Psu complex II), EMD-17640 (https://www.ebi.ac.uk/pdbe/entry/emdb/EMD-17640; ρ^P167L^-ATPγS-Psu complex I) and EMD-17641 (https://www.ebi.ac.uk/pdbe/entry/emdb/EMD-17641; ρ^P167L^-ATPγS-Psu complex II). Structure coordinates have been deposited in the RCSB Protein Data Bank (https://www.rcsb.org) with accession codes 8PEU (https://www.rcsb.org/structure/8PEU; ρ^wt^-ATPγS-Psu complex I), 8PEW (https://www.rcsb.org/structure/8PEW; ρ^wt^-ATPγS-Psu complex II), 8PEX (https://www.rcsb.org/structure/8PEX; ρ^P167L^-ATPγS-Psu complex I) and 8PEY (https://www.rcsb.org/structure/8PEY; ρ^P167L^-ATPγS-Psu complex II). All other data are contained in the manuscript or the Supplementary Information. Source data are provided with this paper. Structure coordinates used in this study are available from the RCSB Protein Data Bank (https://www.rcsb.org) under accession codes 1PV4 (https://www.rcsb.org/structure/1PV4) and 3RX6 (https://www.rcsb.org/structure/3RX6).

## Acknowledgements

We acknowledge the assistance of the core facility BioSupraMol supported by the Deutsche Forschungsgemeinschaft in electron microscopic analyses. This work was supported by grants from the Deutsche Forschungsgemeinschaft (INST 130/1064-1 FUGG to Freie Universität Berlin; GRK 2473 “Bioactive Peptides”, project number 392923329, to M.C.W.; WA 1126/11-1, project number 433623608, to M.C.W.), the Bundesministerium für Bildung und Forschung (ICMR2019-016 to M.C.W.), the Berlin University Alliance (501_BIS-CryoFac to M.C.W.), the Indian Council of Medical Research (AMR/INDO/GER/219/2019-ECD-IIG to R.S.), and the National Institutes of Health (GM067153 to I.A.).

## Author contributions

D.G. produced recombinant proteins and assembled complexes with help by N.S.. T.H. acquired, processed and refined cryoEM data. D.G. built atomic models with the help of M.C.W. and B.L. and refined models with the help of B.L.. D.G. conducted *in vitro* biochemical and biophysical experiments. N.K. cloned constructs for *in vivo* studies and conducted the *in vivo* pull down, bacterial growth and termination assays. B.W. performed bioinformatic analyses. All authors contributed to the analysis of the data and the interpretation of the results. D.G. wrote the original draft with contributions from I.A. and R.S.. M.C.W. revised the manuscript with contributions from D.G., I.A. and R.S.. I.A., R.S. and M.C.W. supervised work in their respective groups and coordinated the collaboration.

## Competing interests

The authors declare no competing interests.

## Extended data

**Extended Data Fig. 1:**
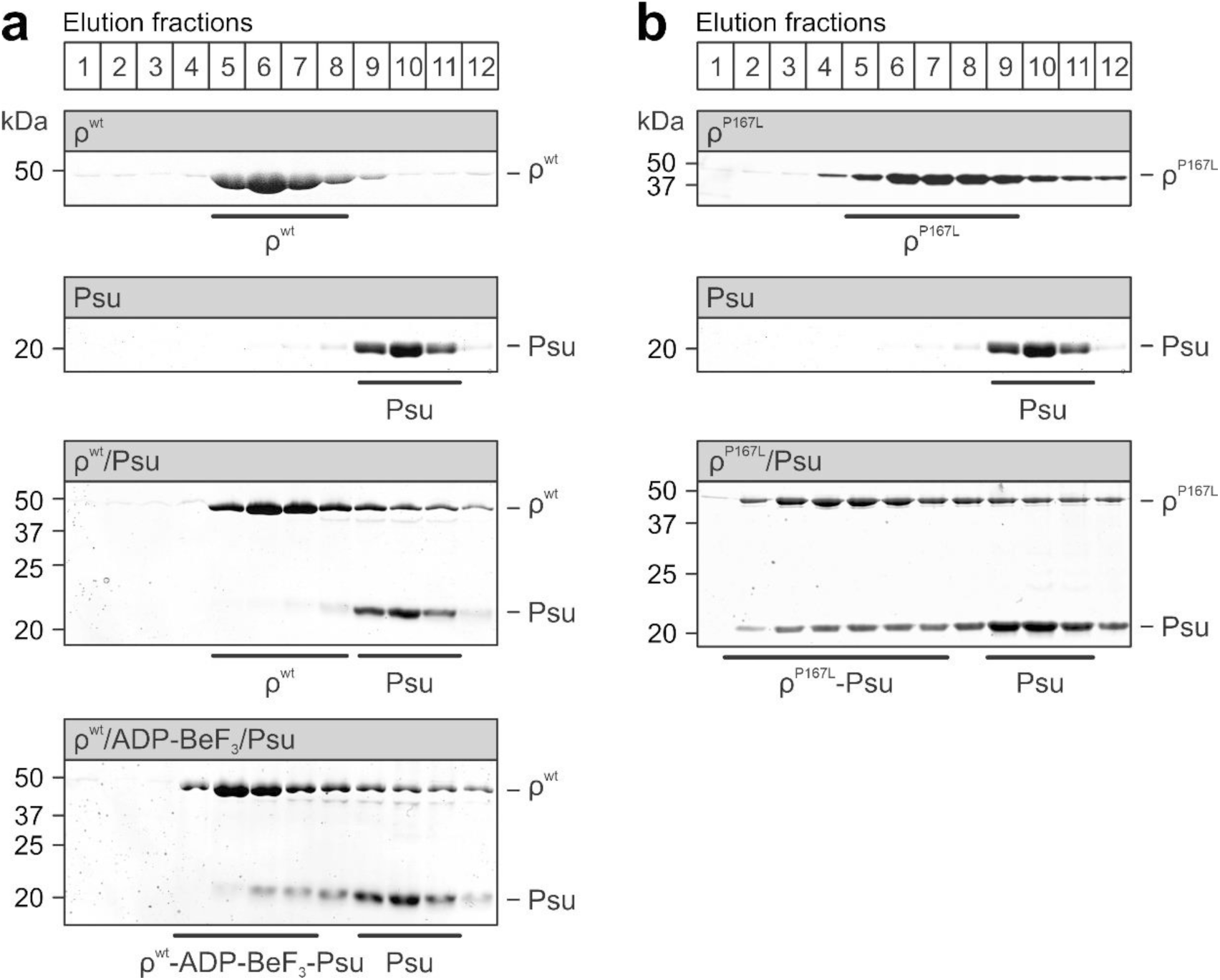
ρ-Psu interaction in solution. **a,b**, SDS-PAGE analyses of analytical SEC elution fractions monitoring the interaction of ρ^wt^ (**a**) or ρ^P167L^ (**b**) with Psu. Elution fractions are indicated at the top. The same elution fractions were analyzed for each run and were aligned below each other. Proteins and protein mixtures analyzed are indicated above each gel. Molecular mass markers are indicated on the left. Protein bands are identified on the right. Fractions containing isolated proteins or complexes are identified below each gel. Experiments were repeated independently at least two times with similar results.

**Extended Data Fig. 2:**
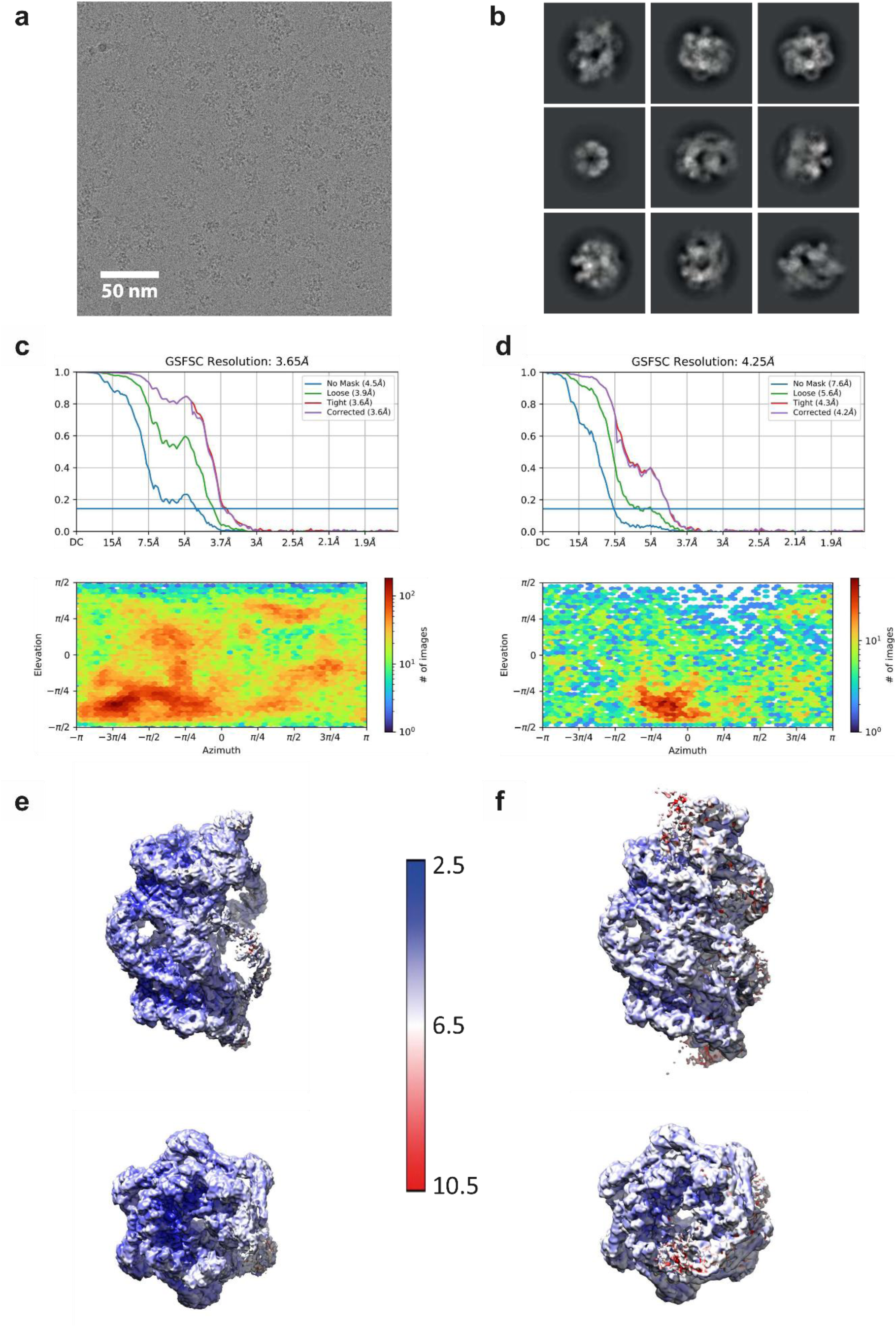
CryoEM/SPA analysis of a ρ^wt^-ATPγS-Psu complexes. **a**, Representative cryoEM micrograph of the ρ^wt^-ATPγS-Psu complexes. Scale bar, 50 nm. 6,266 micrographs were recorded, particle images were picked from 5,986 high-quality micrographs. **b**, 2D class averages of ρ^wt^-ATPγS-Psu particle images after reference-free 2D classification. **c,d**, Upper panels, global resolution estimation for the ρ^wt^-ATPγS-Psu cryoEM reconstructions (complex I, left; complex II, right) by gold-standard Fourier shell correlation (FSC). Blue line, FSC_0.143_. Lower panels, viewing direction distribution plots of the particle images used for the final ρ^wt^-ATPγS-Psu cryoEM reconstructions (complex I, left; complex II, right) as obtained during NU refinement with cryoSPARC. **e,f**, Local resolution estimation as determined with cryoSPARC, ranging from 2.5 Å to 10.5 Å for the ρ^wt^-ATPγS-Psu cryoEM reconstructions (complex I, left; complex II, right).

**Extended Data Fig. 3:**
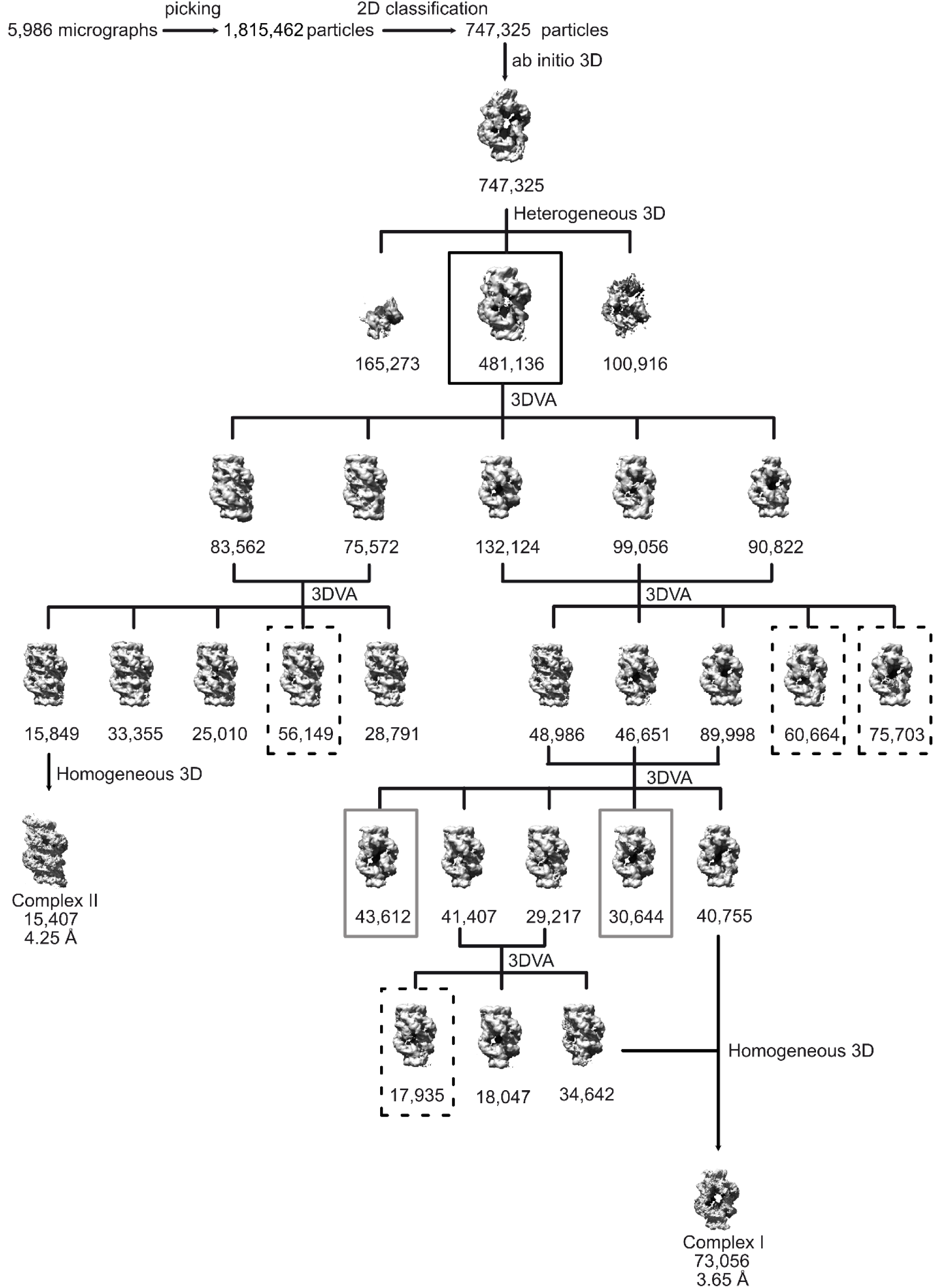
ρ^wt^-ATPγS-Psu cryoEM data refinement. 1,815,462 particle images were picked from 5,986 micrographs and subjected to reference-free 2D classification. 747,325 particle images were selected for iterative cycles of heterogeneous 3D refinement into 3 classes. The best-appearing class, consisting of 481,136 particle images, was selected and subjected to 3D variability analysis (3DVA) generating five clusters. Two of these were combined (left branch) and further classified by 3DVA into five classes, one of which, consisting of 15,849 particles was subjected to homogeneous 3D refinement, which yielded a reconstruction at 4.25 Å resolution (complex II). The residual clusters (right branch) were further classified by iterative 3DVA cycles. Two of the resulting classes appeared virtually identical and were combined for homogeneous refinement yielding a reconstruction at 3.65 Å resolution (complex I). The reconstructions boxed in solid gray boxes represent assemblies with altered rotational orientation of Psu-bridged ρ^wt^ rings, equivalent to ρ^P167L^-ATPγS-Psu complex I. The reconstructions boxed in dashed boxes represent assemblies with intermediate oligomeric states.

**Extended Data Fig. 4:**
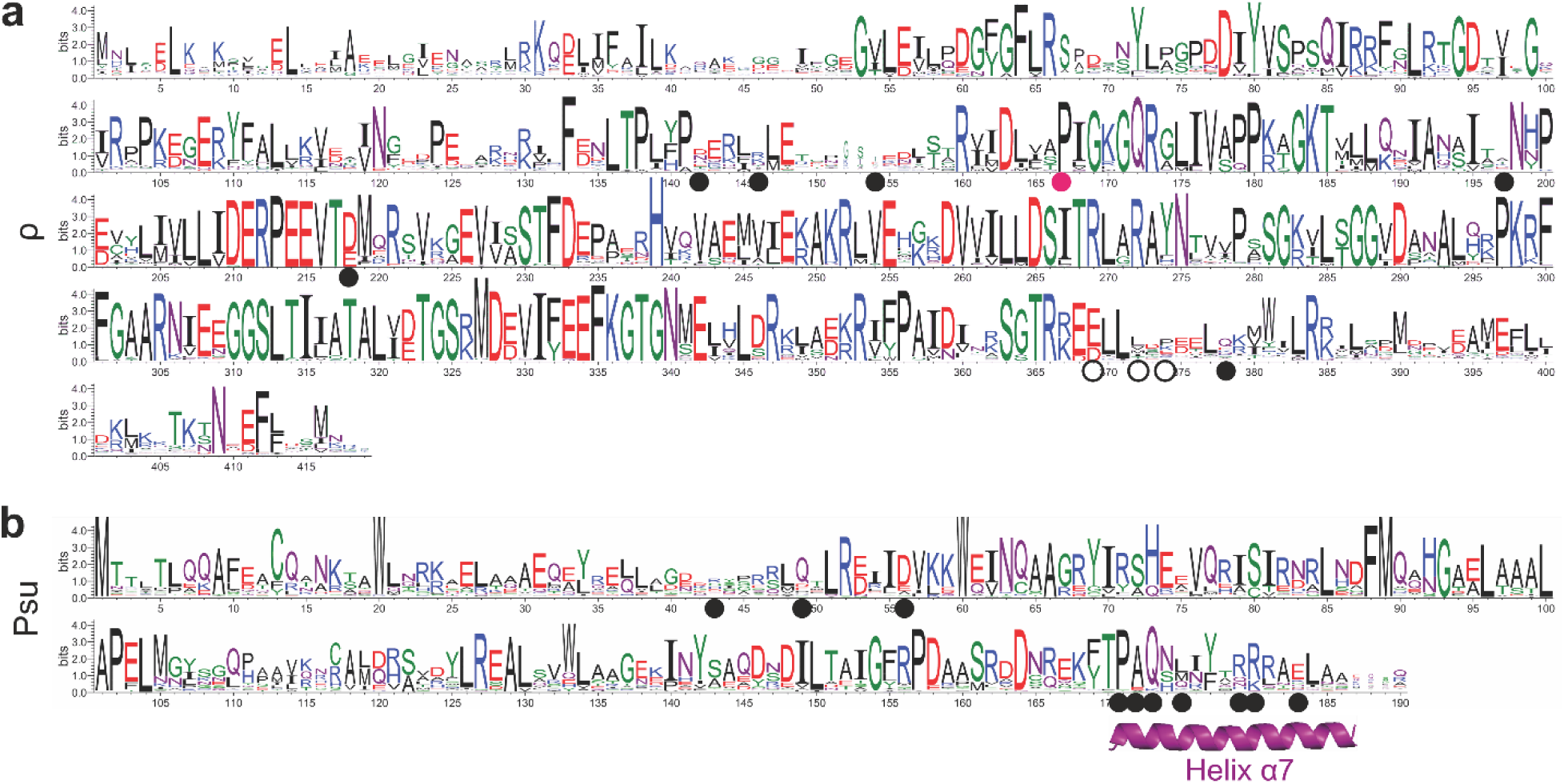
ρ and Psu sequence conservation. **a**, Conservation pattern of ρ based on a multiple sequence alignment of ρ proteins from representatives of the entire Bacteria kingdom (Supplementary Dataset 1A). The universally conserved P167 residue is marked by a magenta sphere. Residues exhibiting side chain interactions between *E. coli* ρ and phage P4 Psu are marked by black spheres; residues exhibiting backbone interactions are marked by white spheres. **b**, Conservation pattern of Psu proteins mined in this study (Supplementary Dataset 1B).

**Extended Data Fig. 5:**
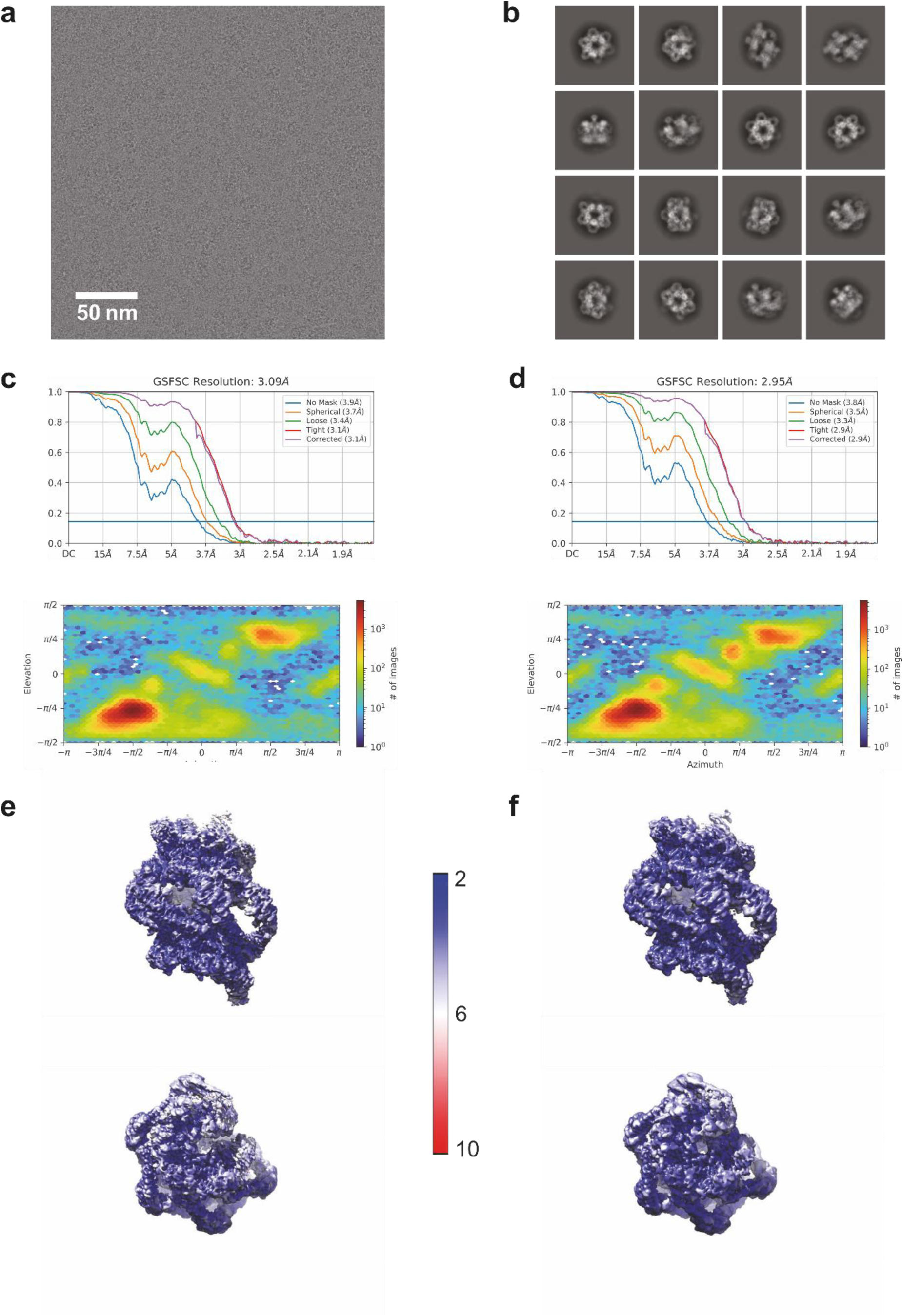
CryoEM/SPA analysis of ρ^P167L^-ATPγS-Psu complexes. **a**, Representative cryoEM micrograph of the ρ^P167L^-ATPγS-Psu complexes. Scale bar, 50 nm. 2,766 micrographs were recorded, particle images were picked from 2,723 high-quality micrographs. **b**, 2D class averages of ρ^P167L^-ATPγS-Psu particle images after reference-free 2D classification. **c,d**, Upper panels, global resolution estimation for the ρ^P167L^-ATPγS-Psu cryoEM reconstructions (complex I, left; complex II, right) by gold-standard Fourier shell correlation (FSC). Blue line, FSC_0.143_. Lower panels, viewing direction distribution plots of the particle images used for the final ρ^P167L^-ATPγS-Psu cryoEM reconstructions (complex I, left; complex II, right) as obtained during NU refinement with cryoSPARC. **e,f**, Local resolution estimation as determined with cryoSPARC, ranging from 2.5 Å to 10 Å for the ρ^P167L^-ATPγS-Psu cryoEM reconstructions (complex I, left; complex II, right).

**Extended Data Fig. 6:**
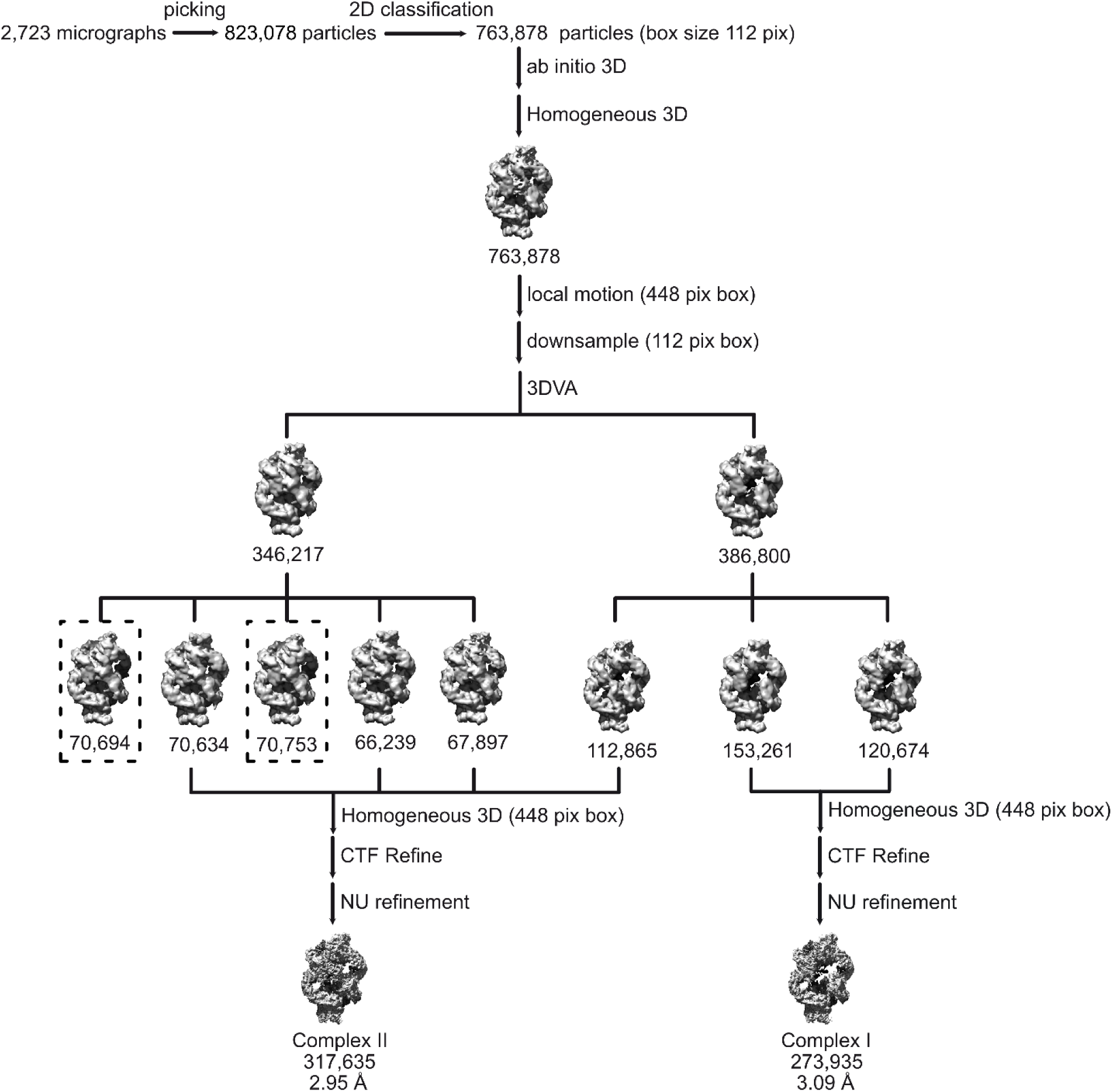
ρ^P167L^-ATPγS-Psu cryoEM data refinement. 823,078 particle images were picked from 2,723 micrographs and subjected to reference-free 2D classification. 763,878 particle images were selected and used for *ab initio* 3D reconstruction, followed by homogeneous refinement. Particles were subjected to local motion correction and re extracted with a box size of 448 px. After down-sampling to a box size of 112 px, 3DVA was applied to separate the dataset into two classes. Each of these was individually subjected to 3DVA, yielding five and three classes, respectively. Similar appearing classes of both branches were combined and homogeneously refined followed by global and local CTF refinement. Final NU refinement yielded reconstructions of 317,635 particle images at 2.95 Å resolution (complex II) and 273,935 particle images at 3.09 Å resolution (complex I). The remaining two classes highlighted by dashed boxes represent additional assemblies.

**Extended Data Fig. 7:**
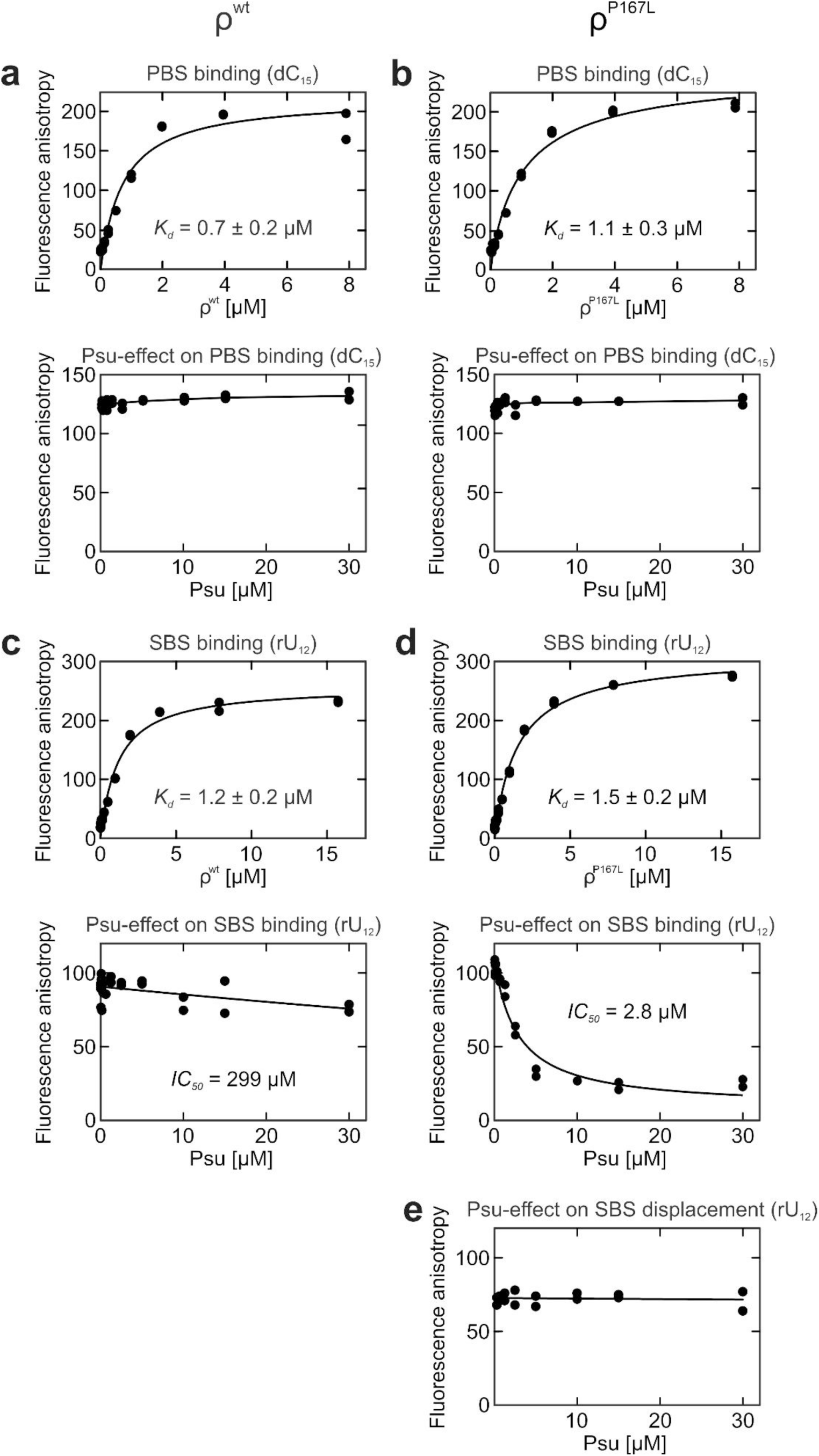
Effects of Psu on nucleic acid binding at the ρ PBS and SBS. **a,b**, Fluorescence anisotropy measurements monitoring the interaction of increasing concentrations of ρ^wt^ (**a**) or ρ^P167L^ (**b**) with dC_15_-FAM at the PBSes (top), or the effect of increasing concentrations of Psu on ρ^wt^ (**a**) or ρ^P167L^ (**b**) dC_15_-FAM binding at the PBSes (bottom). **c,d**, Fluorescence anisotropy measurements monitoring the interaction of increasing concentrations of ρ^wt^ (**c**) or ρ^P167L^ (**d**) with rU_12_-FAM substrate at the SBS (top), or the effect of increasing concentrations of Psu on ρ^wt^ (**c**) or ρ^P167L^ (**d**) rU_12_-FAM binding at the SBS (bottom). **e**, Lack of Psu-mediated release of SBS-bound RNA. Data were recorded as technical duplicates, n = 2. Data in (**a-d**, top) were fit to a single exponential Hill equation for dC_15_-FAM or rU_12_-FAM association at ρ PBSes or SBS. *A* = *A_max_*[protein]^h^/(*K*_d_ ^h^ + [protein]^h^); in which *A_max_* is the fitted maximum of nucleic acid bound; *K_d_* is the dissociation constant; h is the Hill coefficient. Data in (**c,d**, bottom) were fit to an [inhibitor] *vs.* response equation: *A* = *A_min_*+(*A_max_*-*A_min_*)/(1+([inhibitor]/*IC*_50_)); in which *A* is the anisotropy signal at a given concentration of Psu (inhibitor); *A_min_* and *A_max_* are the fitted minimum and maximum of nucleic acid bound.

**Extended Data Fig. 8:**
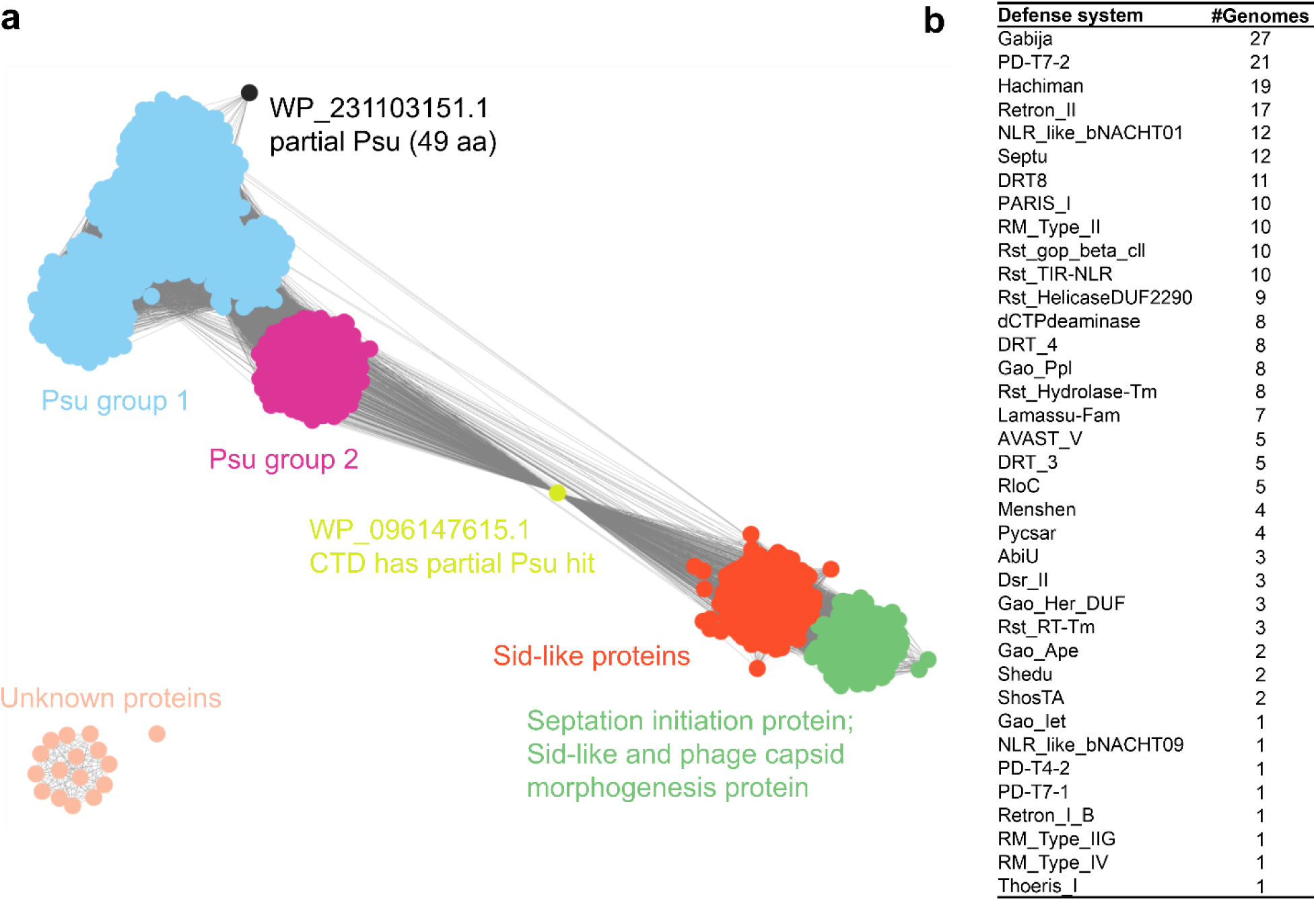
Psu homologs and associated defense systems. **a**, The two Psu groups from the ‘greedy search’. See Methods for details. **b**, A complete count of the DefenseFinder hits.

## Supplementary information

### Supplementary Tables

**Supplementary Table 1:** CryoEM data collection, refinement and validation statistics.

| | $\rho^{\text{wt-ATPyS-Psu}}$ | | $\rho^{\text{P167L-ATPyS-Psu}}$ | |
| --- | --- | --- | --- | --- |
|  | Complex I<br>(PDB 8PEU)<br>(EMDB 17637) | Complex II<br>(PDB 8PEW)<br>(EMDB 17639) | Complex I<br>(PDB 8PEX)<br>(EMDB 17640) | Complex II<br>(PDB 8PEY)<br>(EMDB 17641) |
| Data collection and processing |  |  |  |  |
| Microscope | FEI Titan Krios G3i |  |  |  |
| Voltage [keV] | 300 |  |  |  |
| Camera | Falcon 3EC |  |  |  |
| Magnification (nominal / calibrated) | 96,000 | 96,000 | 96,000 | 96,000 |
| Pixel size at detector [Å/pixel] | 0.832 | 0.832 | 0.832 | 0.832 |
| Total electron exposure [e<sup>-</sup>/Å<sup>2</sup>] | 42 | 42 | 42 | 42 |
| Exposure rate [e<sup>-</sup>/pixel/s] | 0.7 | 0.7 | 0.7 | 0.7 |
| Frames collected during exposure | 33 |  |  |  |
| Defocus range [μm] | 0.80 - 2 |  |  |  |
| Automation software | EPU (version 2.8.1) |  |  |  |
| Micrographs collected | 6,066 | 6,066 | 2,766 | 2,766 |
| Micrographs used | 5,986 | 5,986 | 2,723 | 2,723 |
| Total extracted particles | 1,815,462 | 1,815,462 | 823,078 | 823,078 |
| Final particles | 73,056 | 15,407 | 273,935 | 317,635 |
| Point-group or helical symmetry parameters | C1 | C1 | C1 | C1 |
| Global resolution [Å]; FSC<sup>a</sup><sub>0.143</sub> (unmasked / masked) | 4.5 / 3.6 | 7.6 / 4.2 | 3.9 / 3.1 | 3.8 / 2.9 |
| Local resolution range [Å] | 1.8-30.0 | 3.1-30.0 | 2.2-30.0 | 2.1-30.0 |
| Map sharpening *B* factor [Å<sup>2</sup>] / (*B* factor range) | -82 | -57 | -81 | -83 |
| Map sharpening method | Local *B* factor |  |  |  |
| Refinement package | PHENIX (version 1.20_4459); real.space.refine |  |  |  |
| Model composition |  |  |  |  |
| Non-hydrogen atoms | 57,632 | 83,524 | 54,751 | 55,957 |
| $\rho$ copies / residues | 12 / 39,588 | 18 / 52,588 | 12 / 39,588 | 13 / 41,172 |
| Psu copies / residues | 10 / 14,780 | 16 / 23,468 | 10 / 14,780 | 10 / 14,780 |
| ATPyS molecules | 12 | 16 | 12 | 12 |
| Mg<sup>2+</sup> ions | 12 | 16 | 12 | 12 |
| Model Refinement |  |  |  |  |
| Model-map scores |  |  |  |  |
| CC<sup>b</sup> (mask) | 0.71 | 0.72 | 0.83 | 0.85 |
| CC (volume) | 0.69 | 0.68 | 0.82 | 0.85 |
| Average *B* factors [Å<sup>2</sup>] |  |  |  |  |
| Overall | 162 | 216 | 167 | 150 |
| $\rho$ copies | 162 | 226 | 171 | 149 |
| Psu copies | 162 | 190 | 156 | 153 |
| ATPyS molecules | 202 | 277 | 130 | 129 |
| Mg<sup>2+</sup> ions | 221 | 282 | 113 | 120 |
| Rmsd<sup>c</sup> from ideal values |  |  |  |  |
| Bond lengths [Å] | 0.002 | 0.003 | 0.003 | 0.003 |
| Bond angles [°] | 0.621 | 0.713 | 0.634 | 0.606 |
| Validation<sup>d</sup> |  |  |  |  |
| MolProbity score | 1.72 | 1.96 | 1.15 | 1.69 |
| CaBLAM outliers [%] | 0.9 | 1.0 | 1.1 | 1.2 |
| Clashscore | 14.3 | 18.7 | 13.8 | 13.8 |
| Poor rotamers [%] | 0.0 | 0.0 | 0.0 | 0.0 |
| Cβ deviations | 0 | 0 | 0 | 0 |
| EMRinger score | 0.78 | -0.26 | 1.56 | 1.50 |
| **Ramachandran plot** |  |  |  |  |
| Favored [%] | 97.7 | 96.9 | 98.1 | 97.8 |
| Allowed [%] | 2.3 | 3.1 | 1.9 | 2.2 |
| Outliers [%] | 0.0 | 0.0 | 0.0 | 0.0 |
| **Ramachandran Z-score (rmsd)** |  |  |  |  |
| Whole | 0.78 (0.10) | -0.10 (0.08) | 0.82 (0.10) | 0.75 (0.10) |
| Helix | 1.18 (0.07) | 0.48 (0.07) | 1.28 (0.09) | 1.32 (0.09) |
| Sheet | 0.34 (0.19) | 0.00 (0.16) | 0.25 (0.19) | -0.12 (0.18) |
| Loop | -0.48 (0.13) | -0.84 (0.10) | -0.40 (0.12) | -0.58 (0.13) |

a FSC, Fourier shell correlation
b CC, correlation coefficient
c Rmsd, root-mean-square deviation
d Using MolProbity<sup>91</sup>

**Supplementary Table 2:** Defective Psu and ρ variants.

| **Psu variants defective in $\rho$ binding/inhibition** |  |  |
| --- | --- | --- |
| **Variant** | **Likely effect** | **Reference** |
| L21P | Destabilization of the $\alpha 1/\alpha 2$ coiled-coil | 42 |
| V45F | Local mis-folding, disruption of R43 and D49 interactions with Y197 and R146 of $\rho$ | 16,19 |
| S72L | Destabilization of the $\alpha 1/\alpha 2$ coiled-coil | 42 |
| E56K | Disruption of the interaction with the backbone amide of $\rho$ Q374 | 42 |
| $\Delta 95-98$ | Disruption of Psu dimerization | 37 |
| $\Delta 97-100$ | Disruption of Psu dimerization | |
| P157L | Destabilization of Psu dimer | 42 |
| P157S | Destabilization of Psu dimer | 42 |
| R166C | Effect on helix $\alpha 7$ positioning | 42 |
| R166P | Effect on helix $\alpha 7$ positioning | 42 |
| F169V | Effect on helix $\alpha 7$ positioning through stacking with W60 | 37 |
| ΔCTD10 | Partial deletion of helix $\alpha 7$ interaction interface | 38 |
| ΔCTD20 | Deletion of helix $\alpha 7$ interaction interface | 38 |
| **$\rho$ variants defective in Psu binding/resistant to Psu** |  |  |
| **Variant** | **Likely effect** | **Reference** |
| R144E | Local mis-folding, effect on interaction of R146 with D49 of Psu | 33 |
| $\Delta 144-148$ | Local mis-folding, disruption of interaction with D49 of Psu | 33 |
| R146E | Disruption of interaction with D49 of Psu | 33 |
| E148R | Local mis-folding | 33 |
| R149E | Disruption of stacking with Y197 of $\rho$ to position it for interaction with R43 of Psu | 33 |
| $\Delta 149-153$ | Local mis-folding, effect on the positioning of Y197 of $\rho$ for interaction with R43 of Psu | 33 |
| N151D | Effect on the positioning of Y197 of $\rho$ for interaction with R43 of Psu | 33 |

**Supplementary Table 3:**
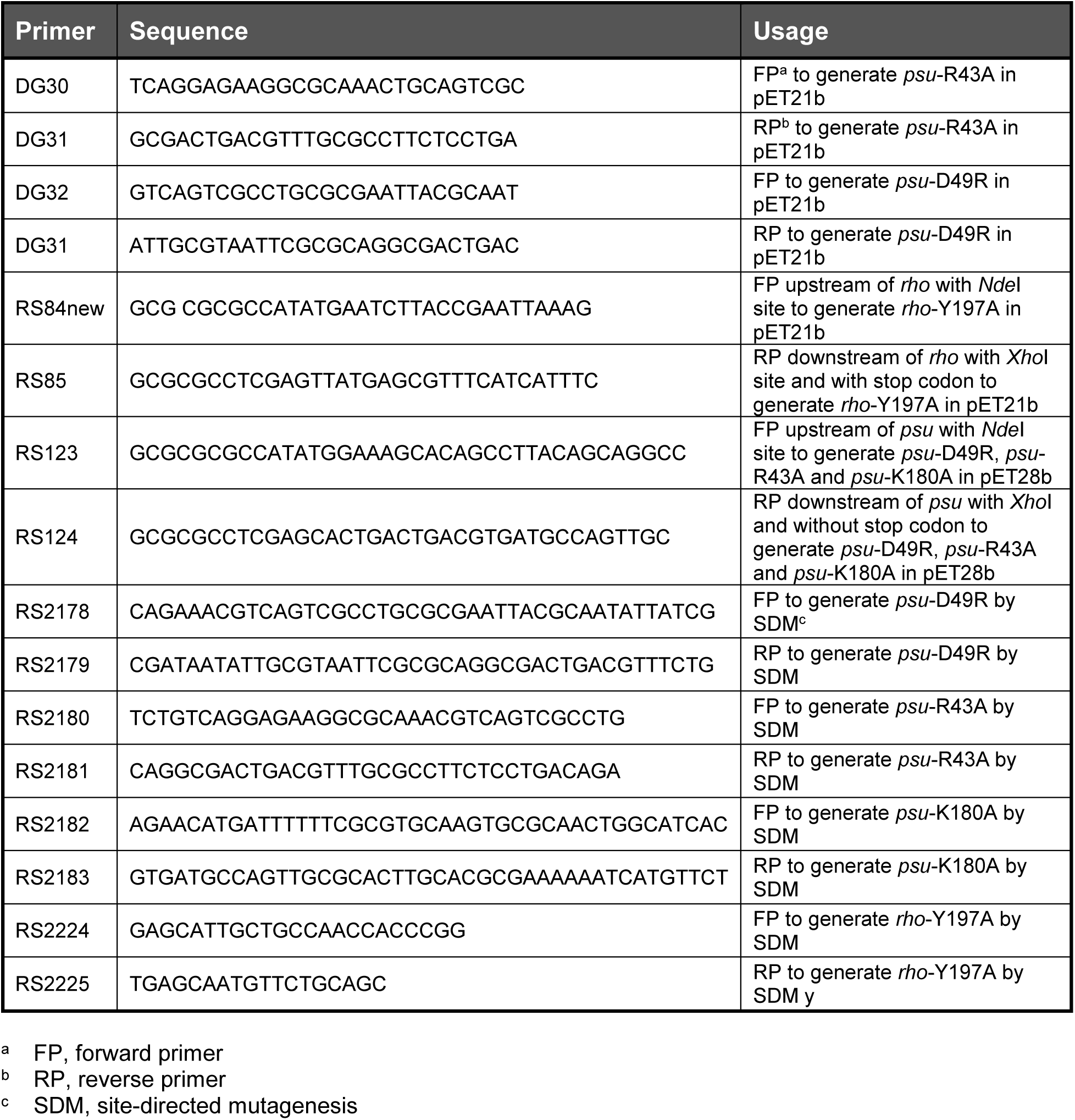
PCR primers (5’ to 3’).

**Supplementary Table 4:** Plasmids and strains.

| Plasmid | Description | Source |
| --- | --- | --- |
| pETM11-*rho* | pETM11 with wt *rho*; no encoded tag; *Kan*<sup>R</sup> | 9 |
| pUC18-*λt<sub>R1</sub>-rut* | pUC19 with the *λt<sub>R1</sub> rut* region cloned at *Xba*I/*Kpn*I sites; *Amp*<sup>R</sup> | 27 |
| pRS455 | pET21b with wt *psu* cloned at *Nde*I/*Xho*I sites; no encoded tag; *Amp*<sup>R</sup> | This study |
| pRS964 | pET21b with *rho*-P167L cloned at *Nde*I/*Xho*I sites; no encoded tag; *Amp*<sup>R</sup> | 33 |
| pRS100 | pET21b with wt *rho* cloned at *Nde*I/*Xho*I sites; no encoded tag; *Amp*<sup>R</sup> | 38 |
| pRS259 | pNL150 with *psu*-F169V; *Cam*<sup>R</sup> | 37 |
| pRS458 | pET28b with wt *psu* cloned at *Nde*I/*Xho*I sites; encoded N-terminal His-tag; *Kan*<sup>R</sup> | 38 |
| pRS555 | pNL150 with *psu*-E56K; *Cam*<sup>R</sup> | 38 |
| pRS624 | pET28b with *psu*-E56K cloned at *Nde*I/*Xho*I sites; encoded N-terminal His-tag; *Kan*<sup>R</sup> | 42 |
| pRS649 | pCL1920 with wt *rho* with its promoter cloned at *Hind*III/*Sac*I sites; *Spec*<sup>R</sup>, *Strep*<sup>R</sup> | 93 |
| pRS1117 | pNL150 with wt *psu* | 37 |
| pRS1199 | RS649 with *rho*-R146E; *Spec*<sup>R</sup>, *Strep*<sup>R</sup> | 33 |
| pRS1219 | pET21b with *rho*-R146E cloned at *Nde*I/*Xho*I site; no encoded tag; *Amp*<sup>R</sup> | 33 |
| pRS2158 | pNL150 with *psu*-D49R; *Cam*<sup>R</sup> | This study |
| pRS2159 | pNL150 with *psu*-K180A; *Cam*<sup>R</sup> | This study |
| pRS2160 | pNL150 with *psu*-R43A; *Cam*<sup>R</sup> | This study |
| pRS2194 | RS649 with *rho*-Y197A; *Spec*<sup>R</sup>, *Strep*<sup>R</sup> | This study |
| pRS2236 | pET21b with Y197A *rho* cloned at *Nde*I/*Xho*I site, Non-His-tag; *Amp*<sup>R</sup> | This study |
| pRS2237 | pET28b with R43A *psu* cloned at *Nde*I/*Xho*I site, His-tag at N-terminal; *Kan*<sup>R</sup> | This study |
| pRS2238 | pET28b with D49R *psu* cloned at *Nde*I/*Xho*I site, His-tag at N-terminal; *Kan*<sup>R</sup> | This study |
| pRS2239 | pET28b with K180A *psu* cloned at *Nde*I/*Xho*I site, His-tag at N-terminal; *Kan*<sup>R</sup> | This study |
| Strain | Genotype | Source |
| RS1263 | *E. coli* MG1655; K-12 | Lab stock |
| RS1309 | *E. coli* MG1655Δ*rho*Δ*rac*, with shelter plasmid pHYD1201 (*Amp*<sup>R</sup>) | 107 |
| RS2047 | MG1655$\Delta rac\Delta lac$, λRS45 lysogen carrying $P_{lac}$-$\lambda t_{R1}$-$lacZYA$ | 75 |

## Notes

### Competing Interest Statement

The authors have declared no competing interest.

